# Mouse paralaminar amygdala excitatory neurons migrate and mature during adolescence

**DOI:** 10.1101/2022.09.23.509244

**Authors:** P.J. Alderman, D. Saxon, L.I. Torrijos-Saiz, M. Sharief, S.W. Biagiotti, C.E. Page, A. Melamed, C.T. Kuo, J.M. Garcia-Verdugo, V. Herranz-Pérez, J.G. Corbin, S.F. Sorrells

**Affiliations:** Department of Neuroscience, University of Pittsburgh, Pittsburgh PA, 15260; Center for Neuroscience Research, Children’s Research Institute, Children’s National Hospital, Washington, DC, USA; Interdisciplinary Program in Neuroscience, Georgetown University Medical Center, Washington, DC, United States; Laboratory of Comparative Neurobiology. Institute Cavanilles. University of Valencia, CIBERNED, Valencia, 46980, Spain; Department of Cell Biology, Functional Biology and Physical Anthropology, University of Valencia, Burjassot, 46100, Spain; Department of Cell Biology, Duke University School of Medicine, Durham, NC 27710. United States

**Keywords:** Migration, amygdala, neurogenesis, maturation, endopiriform, intercalated

## Abstract

The human amygdala paralaminar nucleus (PL) contains immature excitatory neurons that exhibit protracted maturation into adolescence; however, whether a similar population exists in mice is unknown. We discovered a previously undescribed region with immature doublecortin (Dcx)^+^ excitatory neurons adjacent to the mouse basolateral amygdala, and similar to humans, these neurons mature during adolescence and are distinct from adjacent intercalated cells. Despite their immature features, these neurons are born during embryogenesis, populate the mouse PL prior to birth, and remain in an immature stage of development until adolescence. In the postnatal brain, a subpopulation of these excitatory neurons surprisingly migrate into the neighboring endopiriform cortex, peaking between P21–P28. In humans, cells with the molecular identity of mouse PL neurons populate the PL as early as 18 gestational weeks, and also exhibit migratory morphology into adolescence (13 years). The finding of a similar region in both mice and humans suggests a potentially conserved cellular mechanism for neuron recruitment and migration during adolescence, a key time period for amygdala circuit maturation and behavioral changes.

## Introduction

The amygdala is a brain region centrally involved in emotional learning and salience discrimination^1, 2^, processes that are refined in childhood and adolescence^3–5^. From infancy into adolescence, the human amygdala continues to grow significantly in size and neuron number^6–10^. Part of this growth arises from the continued maturation of a unique population of doublecortin (DCX)-expressing immature excitatory neurons present in the amygdala paralaminar nucleus (PL)^6,7,11,12^. The recruitment of these immature neurons into established circuits is proposed to influence circuit plasticity^13–16^, however little is known about the cellular mechanisms underlying this unique form of neuron development.

In humans, the PL was first anatomically distinguished by analysis of simple histological preparations^17^. Human PL neurons express the immature neuron markers DCX^6,7^ and polysialylated neural cell adhesion molecule (PSA-NCAM)^7,11^ and have small nuclei with compacted chromatin and few processes^7^. These immature neurons develop into mature excitatory neurons along a delayed timeline, apparently arrested relative to surrounding neurons. From infancy into childhood and adolescence these immature neurons grow in size and complexity as they downregulate DCX expression, though a subpopulation remains immature as old as 77 years^6,7^. In both humans and non-human primates, the PL is a prominent, densely packed region adjacent to the basolateral amygdala (BLA)^12,18–20^, consisting of approximately 3 million, or ~20% of the neurons, in the adult human amygdala^21^. A similar, but smaller collection of immature neurons has been observed adjacent to the BLA in other mammals^22^ including sheep^23^, cats^11,24^, rabbits^25^, bats^26^, and tree shrews^27^. Whether an experimentally tractable system like the laboratory mouse has a molecularly and developmentally similar population of immature neurons in the adolescent amygdala has not yet been determined^12^.

In humans and other primates, a subset of PL neurons display migratory morphology suggesting that these cells maintain a capacity to migrate during adolescence, however this has not been directly investigated. Postnatal migration of inhibitory interneurons is prominent in the adult rostral migratory stream (RMS)^28^ and early postnatal mouse forebrain^29^. In contrast, excitatory neuron migration is more rare in the adult brain, with local migration of dentate granule neurons^30^ and longer excitatory neuron migration in the avian brain^31^. Since PL neurons are excitatory, their migration into nearby brain regions would be remarkable, and could contribute to observed increases in neuron number in some amygdala regions^6,7,32^. The neuronal identity, migratory potential, and destinations of immature PL neurons have not been linked in any species, but a rodent model would permit this in vivo.

The presence of immature neurons in the amygdala also raises the question of when these cells are born^33–35^. The ages when the PL forms in humans are not clear^36^, however, our previous investigations suggested the region might form as early as 22 weeks of gestation^7^. The number of dividing progenitor cells in the human PL drops sharply between birth and 5 months, much earlier than the gradual decline in immature neurons, suggesting that the majority of immature PL neurons are born during embryogenesis^7,8^. Interestingly, birthdating studies that administered 5-bromo-2-deoxyuridine (BrdU) to adult animals found BrdU co-labeled cells expressing DCX^34^ or NEUN^33^ in the non-human primate amygdala and rodent amygdala^37,38^ suggesting that some amygdala neurons could be born in adults. These findings require further investigation because dividing oligodendrocyte progenitors (OPCs) can express Dcx^39–41^, are abundant in the amygdala^42,43^, and are often closely associated with neurons^39,44^. Whether immature neurons in the PL arise from postnatal neurogenesis remains unknown in any species where these cells have been observed.

We identified a region adjacent to the mouse BLA that we refer to as the mouse PL based on its morphological, molecular, and developmental similarities to the immature excitatory neurons in the human PL. We reveal the cellular mechanisms of its development including local maturation, migration, postnatal and embryonic neurogenesis, and cell death. This study uncovers a conserved developmental mechanism during adolescence in mice and humans whereby excitatory neurons continue to grow and migrate within established circuits. In humans where the PL contains 15-20% of amygdala neurons, changes in this immature population are linked to autism spectrum disorder (ASD)^6^, epilepsy^45^, and/or major depressive disorder (MDD)^46^ and occur during key adolescent ages for neurotypical behavioral development.

## Results

### Immature neurons are present in the postnatal mouse PL

To investigate whether immature neurons are present in the mouse amygdala, we examined Dcx expression at P21 (**Fig. 1, Extended Data Fig. 1,2**). We observed a dense nucleus of Dcx^+^NeuN^+^ and Dcx^+^Psa-Ncam^+^ cells anterior and ventral to the BLA that was most evident in sagittal planes (**Fig. 1a, Extended Data Fig. 1a,b,d,e**). This region was closely associated, but distinct from the Dcx^−^ amygdala intercalated cell clusters (ITCs) (**Fig. 1a–c, Extended Data Figs. 1b, 2b–e, 3a–d**). In coronal sections, the nucleus appears as a small cluster ventrolateral to the striatum at the ventral terminus of the external capsule (**Extended Data Fig. 1c, 2a–e**) and the adjacent ventral endopiriform cortex (vEN).

**Figure 1:**
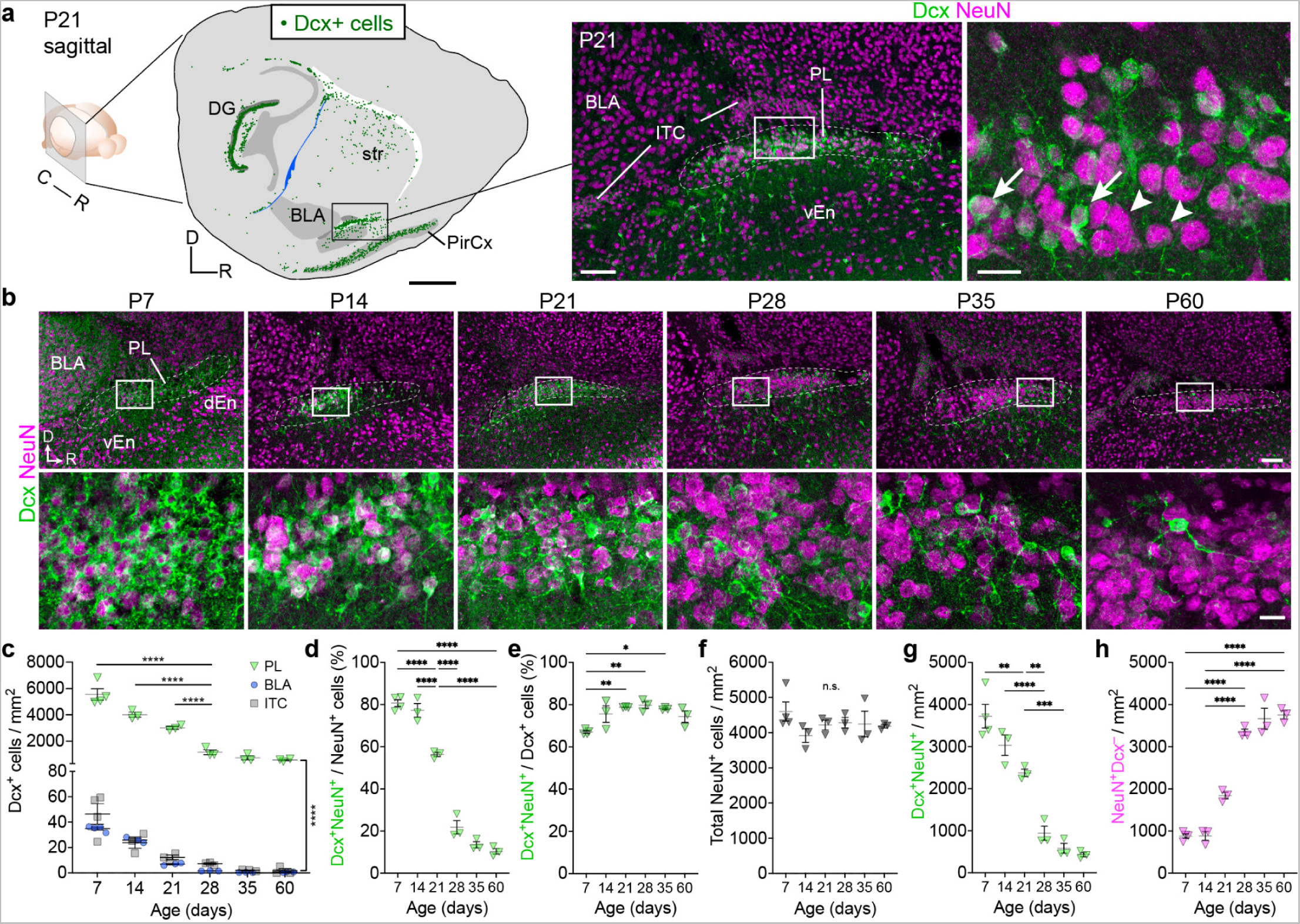
A region with immature neurons in the juvenile mouse amygdala. **a,** Sagittal P21 mouse brain section with green dots indicating Dcx^+^ cells. Immunostaining of the boxed region in this section shows Dcx^+^NeuN^+^ immature neurons (arrows) and Dcx^−^NeuN^+^ mature neurons (arrowheads) located anterior and ventral to the BLA and ITCs in the mouse PL. **b,** Age series of immunostaining for Dcx and NeuN in this region between P7 and P60. **c,** Density of Dcx^+^ cells within the ITCs, BLA, and PL between P7 and P60. **d,** Percentage of NeuN^+^ cells in the PL that are Dcx^+^ between P7 and P60. **e,** Percentage of Dcx^+^ cells in the PL that are NeuN^+^ between P7 and P60. **f–h,** Density of all NeuN^+^ cells (**f**) in the PL and the subpopulations that are Dcx^+^NeuN^+^ (**g**) or Dcx^−^NeuN^+^ (**h**). * = p < 0.05, ** = p < 0.01, *** = p < 0.001, **** = p < 0.0001. 1-way (**d–h**) or 2-way (**c**) ANOVA with Holm-Sidak’s post-hoc test. Scale bars = 1 mm (**a**, left), 100 µm (**a** middle, **b** top row), 20 µm (**a** right, **b** bottom row). DG: dentate gyrus; str: striatum; BLA: basolateral amygdala; PirCx: piriform cortex; ITC: intercalated cell clusters; PL: paralaminar nucleus; vEn: ventral endopiriform cortex; dEn: dorsal endopiriform cortex.

In humans, immature PL neurons develop between birth and adolescence^6,7^, so we next investigated the Dcx^+^ cells in mice across an age-series (P7-60). For this study we considered Dcx^+^NeuN^+^ cells to be immature neurons and Dcx^−^NeuN^+^ cells to be mature neurons. At P7, the PL contained a dense cluster of cells with a meshwork of Dcx labeling that was brighter than surrounding regions (**Fig. 1b**). At P14 and P21, Dcx^+^NeuN^+^ cells continued to be visible within the PL and there was a sharp decline in their abundance between P21 and P28, and a smaller decrease between P35 and P60. We quantified these changes compared to the neighboring BLA and ITCs and found that the PL had significantly higher Dcx^+^ cell density at all ages between P7 to P60 (**Fig. 1c**). Across these ages, the percentage of total PL NeuN^+^ cells co-expressing Dcx significantly declined (**Fig. 1d**), while the percentage of Dcx^+^ cells co-expressing NeuN slightly increased (**Fig. 1e**). The total density of NeuN^+^ cells in the PL remained constant as the Dcx^+^NeuN^+^ immature neuron density significantly decreased and the Dcx^−^NeuN^+^ mature neuron density significantly increased with age (**Fig. 1f–h**). Together, these results suggest that, similar to the human PL, this region in mice contains a high density of Dcx^+^ cells at ages corresponding to human late gestation^47,48^ and continuing into infancy (P7-P14), and that these cells sharply decline during mouse adolescent ages (P21-P35). To next investigate the dynamics of molecular marker expression in this region, we conducted an extensive analysis using markers known to label different embryonic and post natal telencephalic populations.

### The PL contains a high density of Tbr1^+^ excitatory neurons

Immature neurons in the human PL express T brain 1 (TBR1), a marker of cortical excitatory neurons and chicken ovalbumin upstream promoter II (COUPTFII), a transcription factor expressed broadly in the amygdala^7^. Our prior single nuclei RNA-sequencing studies further identified *BCL11B*/*CTIP2* as a putative PL marker in humans^7^. We assessed FoxP2 and Tshz1 expression, which label the ITCs^49–51^, the dense GABAergic cell clusters adjacent to the BLA, and SatB2, a marker of the vEN^52–54^. We also examined expression of the pallial marker Pax6^55^, and Sp8, a marker of migratory interneurons in the RMS^56^.

From this analysis we observed a dense population of PL cells co-expressing Dcx and Tbr1, CoupTFII, or Ctip2, with very few Dcx^+^ cells co-expressing FoxP2, SatB2, Pax6, or Sp8 at P3, P7, P21, P60, and P120 (**Fig. 2a, Extended Data 3a–e**). Tbr1 was consistently highly expressed (~80%) by both the immature (Dcx^+^NeuN^+^) and mature (Dcx^−^NeuN^+^) PL neurons (**Fig. 2b,c**). The percentage (**Fig. 2d**) and density (**Fig. 2e**) of Tbr1^+^NeuN^+^ cells co-expressing Dcx decreased significantly with age while the density of mature (Dcx^−^NeuN^+^) Tbr1^+^ cells increased (**Fig. 2f**); however, total NeuN^+^Tbr1^+^ density remained constant (**Fig. 2g**). CoupTFII was also expressed in most (~75%) immature and mature PL neurons at P21, with a significant decrease in co-expression at P60 (**Fig. 2h,i**). Between P3 and P120, the PL contained dense collections of NeuN^+^Tbr1^+^ cells or NeuN^+^CoupTFII^+^ cells, but few NeuN^+^FoxP2^+^ cells or NeuN^+^SatB2^+^ cells (**Fig. 2j, Extended Data Fig. 3e**).

**Figure 2:**
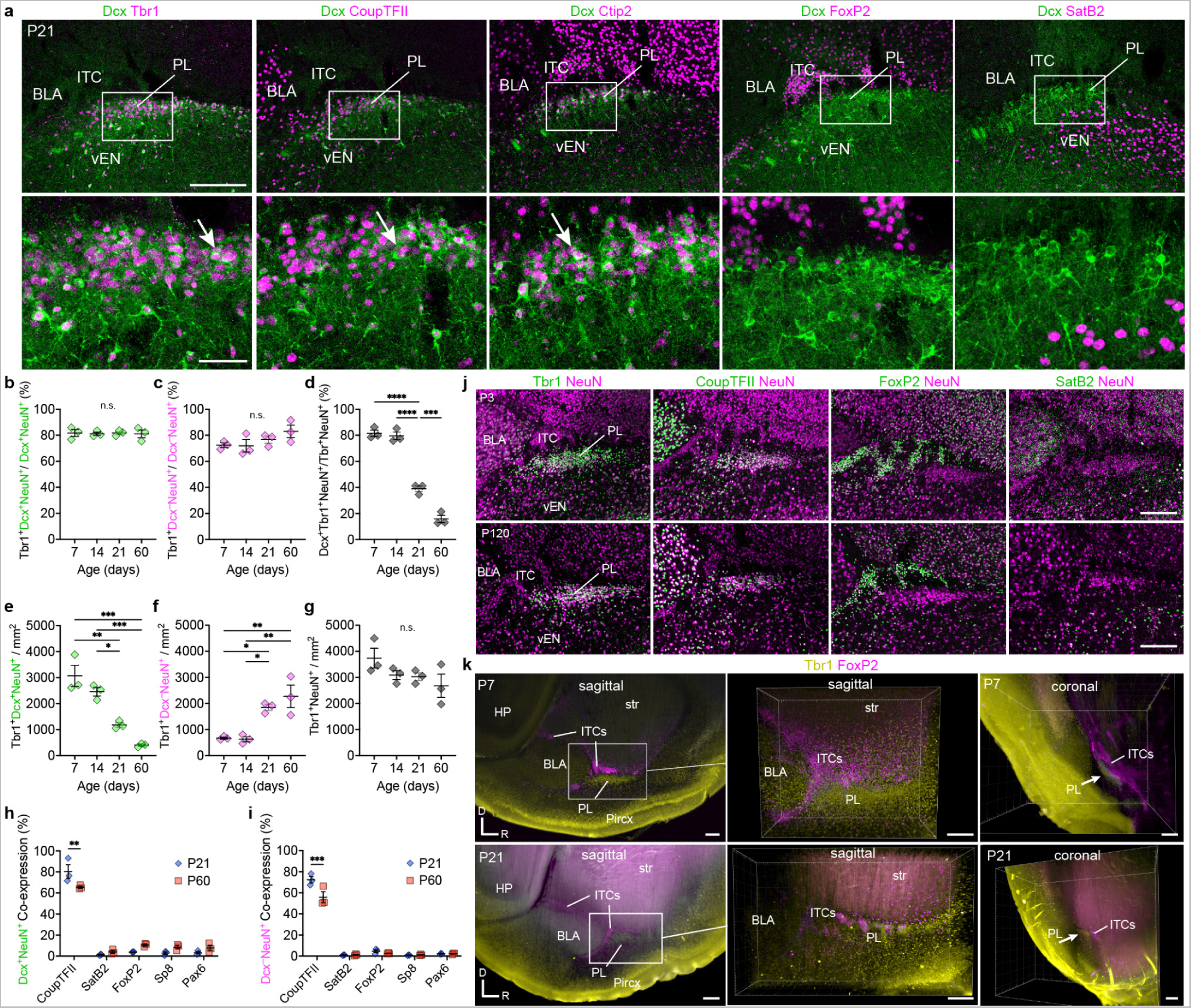
Transcription factor expression pattern within the PL and surrounding regions. **a,** Level-matched sections of the P21 mouse PL co-immunostained for Dcx and either Tbr1, CoupTFII, Ctip2 which label Dcx^+^ cells in the PL (arrows) or the ITC marker FoxP2, or the vEN marker SatB2. **b,c,** Percentage of Dcx^+^NeuN^+^ cells (**b**), or Dcx–NeuN^+^ cells (**c**) co-expressing Tbr1 between P7 and P60. **d,** Percentage of Tbr1^+^NeuN^+^ cells co-expressing Dcx between P7 and P60. **e–g,** Density of Tbr1^+^Dcx^+^NeuN^+^ cells (**e**), Tbr1^+^Dcx^−^NeuN^+^ cells (**f**), or all Tbr1^+^NeuN^+^ cells (**g**) in the PL between P7 and P60. **h,i,** Percentage of Dcx^+^NeuN^+^ cells (**h**) or Dcx–NeuN^+^ cells (**i**) in the PL that co-express each indicated transcription factor at P21 or P60. **j,** Level-matched sections of the P3 and P120 mouse PL immunostained for NeuN together with the PL markers Tbr1 or CoupTFII, and the ITC marker FoxP2, and the vEN marker SatB2. **k,** Three-dimensional imaging of iDISCO cleared brains at P7 and P21 stained for Tbr1 and FoxP2, showing the close proximity of these distinct cell types in the region rostral to the BLA. * = p < 0.05, ** = p < 0.01, *** = p < 0.001, **** = p < 0.0001. 1-way ANOVA with Holm-Sidak’s post-hoc test. Scale bars = 200 µm (**a** top, **j, k**), 50 µm (**a** bottom). BLA: basolateral amygdala; HP: hippocampus; ITC: intercalated cell clusters; PL: paralaminar nucleus; vEn: ventral endopiriform cortex.

We next investigated the anatomical and molecular relationship of neighboring regions to the PL. FoxP2^+^ cells in the ITCs are in close proximity to the PL (**Fig. 2a,j, Extended Data Fig. 2d, 3e**), a relationship most clearly visualized with lightsheet microscopy (**Fig. 2k, Supplementary Movies 1 and 2**). Our analysis also revealed limited overlap (~13%) at P7 of the PL Dcx^+^ cells and Tshz1, which primarily marks dorsal lateral ganglionic eminence (dLGE)-derived cells that populate the ITCs^49–51^. This fraction significantly decreased with age (**Extended Data Fig. 3f–h**). At P21 we also observed a field of Dcx^+^SatB2^−^ cells extending from the PL into the SatB2^+^ vEN (**Fig. 2a,j, Extended Data Fig. 1b–e, 3e**). Together these observations reveal that both immature and mature neurons in the mouse PL are marked by the same regional identity transcription factors as the human PL, and are excitatory, unlike the neighboring ITCs.

### Local PL maturation occurs alongside migration into the vEN

To investigate whether the decrease in PL Dcx^+^ cells and their emanation into the vEN was associated with neuron migration, we next estimated total cell numbers in the PL and vEN between P7 and P60. We conducted unbiased stereology using the optical fractionator probe (**Supplementary Table 1**) to estimate the total population of Dcx^+^ cells (**Fig. 3a–d**). The PL Dcx^+^ cell population decreased by 21.3% from P7–P21, and decreased more rapidly by 63.6% from P21–P28 reaching a plateau between P35–P60. Interestingly, there were still approximately 2,000 Dcx^+^ cells in the PL at P60. The vEN had overall lower numbers of Dcx^+^ cells than the PL; however, at P21, there was a sharp 3-fold increase in Dcx^+^ cells in the vEN. The vEN Dcx^+^ cell population remained elevated at P28 and P35, before falling by P60. The sharp decrease in Dcx^+^ cells in the PL and corresponding increase in the vEN at the same age (P21), suggested the intriguing possibility that a subset of these Dcx^+^ cells might be migrating from the PL to the vEN.

**Figure 3:**
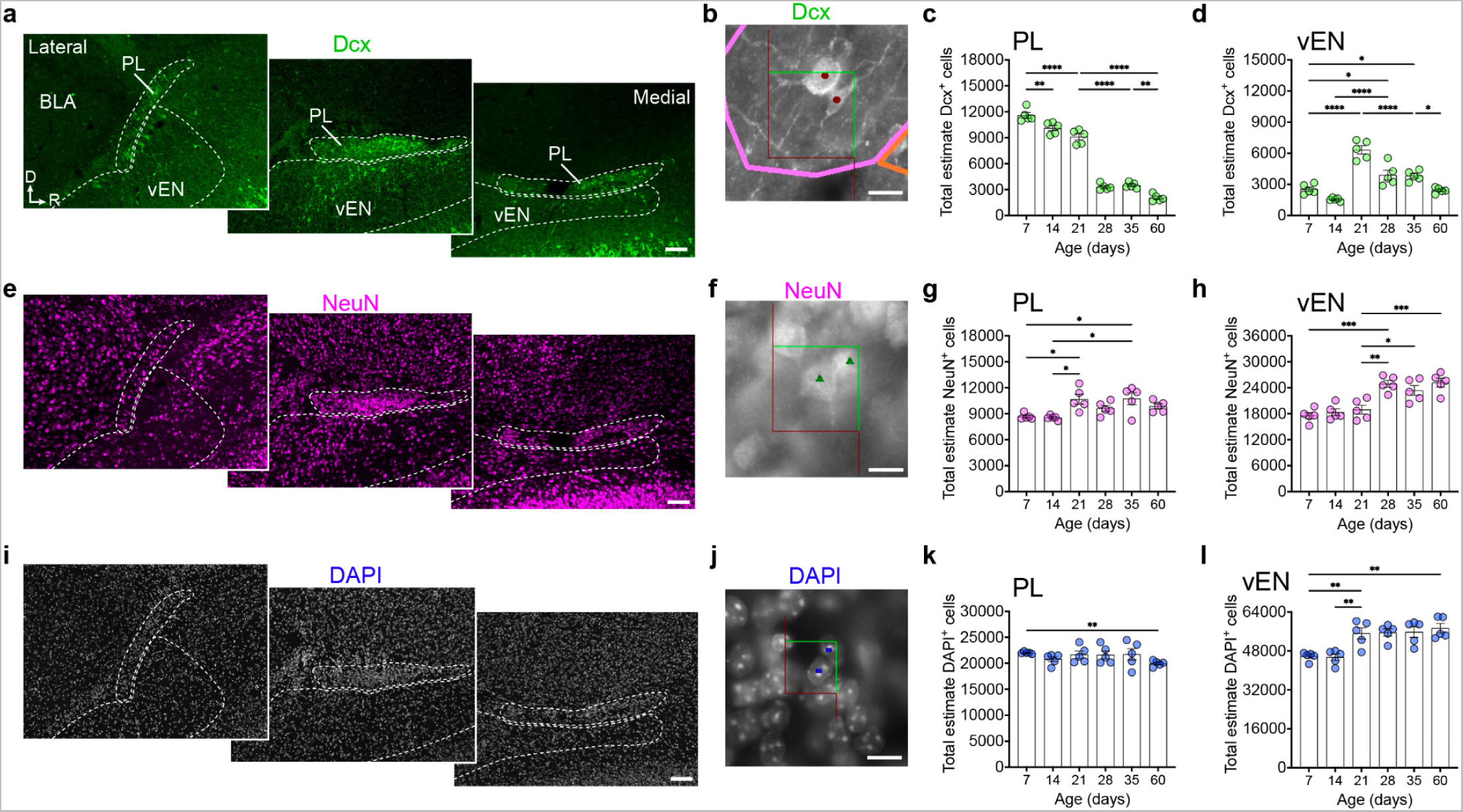
Stereological estimates of total cell numbers in the PL and vEN. Stereological measurements of Dcx^+^ cells (**a–d**), NeuN^+^ cells (**e–h**), and DAPI^+^ nuclei (**i–l**) in the PL and vEN between P7 and P60. Sagittal sections indicating the boundaries traced for the PL across 3 representative medial-lateral levels stained for Dcx (**a**), NeuN (**e**), and DAPI (**i**). Boundaries were established using multiple markers across ages (see Methods for details). The optical fractionator frame size and representative counted objects are shown for Dcx (**b**), NeuN (**f**), and DAPI (**j**). Estimates of total cell populations for Dcx^+^ cells in the PL (**c**) and vEN (**d**), for NeuN^+^ cells in the PL (**g**) and vEN (**h**), and for DAPI^+^ cells in the PL (**k**) and vEN (**l**) between P7 and P60. * = p < 0.05, ** = p < 0.01, *** = p < 0.001, **** = p < 0.0001, 1-way ANOVA with Holm-Sidak’s post-hoc test. Scale bars = 100 µm (**a,e,i**), 10 µm (**b,f,j**). BLA: basolateral amygdala; PL: paralaminar nucleus; vEn: ventral endopiriform cortex.

In the same sections we next estimated the total number of NeuN^+^ cells in each region (**Fig. 3e–h**). In the PL, the total neuronal population was approximately 9,000 neurons at P7, and increased by about 2,000 cells at P21, after which point it remained constant. In the vEN, the total estimate of NeuN^+^ cells was also relatively constant between P7–P21 until P21–P28 when the total number of NeuN^+^ cells in the vEN increased by approximately 5,000. These results supported the possibility that a subset of PL neurons mature locally because NeuN expression is upregulated as immature neurons mature, and a higher percentage of PL Dcx^+^ cells express NeuN with age (**Fig. 1e**). These results also further suggested that a subset of PL Dcx^+^ cells migrated into the vEN because their decrease was much greater than the overall increase in PL NeuN^+^ cells. There was also an increase in vEN NeuN^+^ cells at P28, one week after the appearance of many Dcx^+^ cells in the vEN.

Although total cell number is influenced by postnatal gliogenesis, we next quantified total DAPI^+^ nuclei in each region (**Fig. 3i–l**). The total number of DAPI^+^ nuclei in the PL was approximately twice the number of PL neurons, and exhibited a slight overall decrease with age, dropping by around 2,000 nuclei between P7 and P60. In the vEN, the total DAPI^+^ nuclei remained relatively constant between P7 and P14 before increasing sharply at P21. Together, these quantifications indicated that, coincident with the sharp Dcx^+^ cell decrease in the PL at P21, both Dcx^+^ and DAPI^+^ nuclei increased in the vEN, followed over the next week by an increase in NeuN^+^ cells there. These changes argued that (1) PL Dcx^+^ cells either mature locally within the PL and downregulate Dcx expression, or (2) they remain in the PL where they retain Dcx expression for months, or (3) they migrate into the vEN. It is also possible that cell birth/death could contribute to observed changes in total cell population dynamics. To determine which of these outcomes occur in the PL, we next looked for direct evidence of PL local maturation and/or Dcx expression retention, neurons with migratory properties, and neuron birth or death.

### A subset of immature PL neurons downregulate Dcx and others express it protractedly

The decline in Dcx^+^ cells in the PL suggests that a subpopulation downregulate Dcx expression as they mature. Interestingly, the total Dcx^+^ cell population did not drop to zero by P60, suggesting that a subset (~20%) either retain Dcx expression for months after they are generated, or are newly-born. To study these populations, we used Dcx-CreER^TM^ mice crossed to a ROSA-LSL-tdTomato (tdT) reporter to permanently label Dcx-expressing cells. These mice had expected tamoxifen (TMX)-induced tdT recombination in the rostral migratory stream (RMS), ventricular-subventricular zone (V-SVZ) and olfactory bulb (**Extended Data Fig. 4a,b**) as well as the PL and piriform cortex. We administered TMX to induce recombination immediately prior to the sharp decrease in Dcx cells (once daily, P21–P23, **Fig. 4a–d**), or after the decrease (once daily, P28–P30 **Fig. 4e–g**). We analyzed brains 2, 5, or 12 days after each labeling period and observed simple and complex tdT^+^Dcx^+^Tbr1^+^ recombined cells in the PL, vEN, and piriform cortex from each treatment (**Fig. 4a–g, Extended Data Fig. 4c–f**). In the PL, the percentage of tdT^+^ cells co-expressing Dcx decreased with increasing time post-TMX in the P21–P23 labeling mice; however, this same decrease did not happen from P28–P30 labeling and instead remained significantly elevated at the latest time examined (**Fig. 4h**). These observations indicate that after P21, a subset of PL neurons downregulate Dcx and mature in the PL, and after P28 a subpopulation retains Dcx expression.

**Figure 4:**
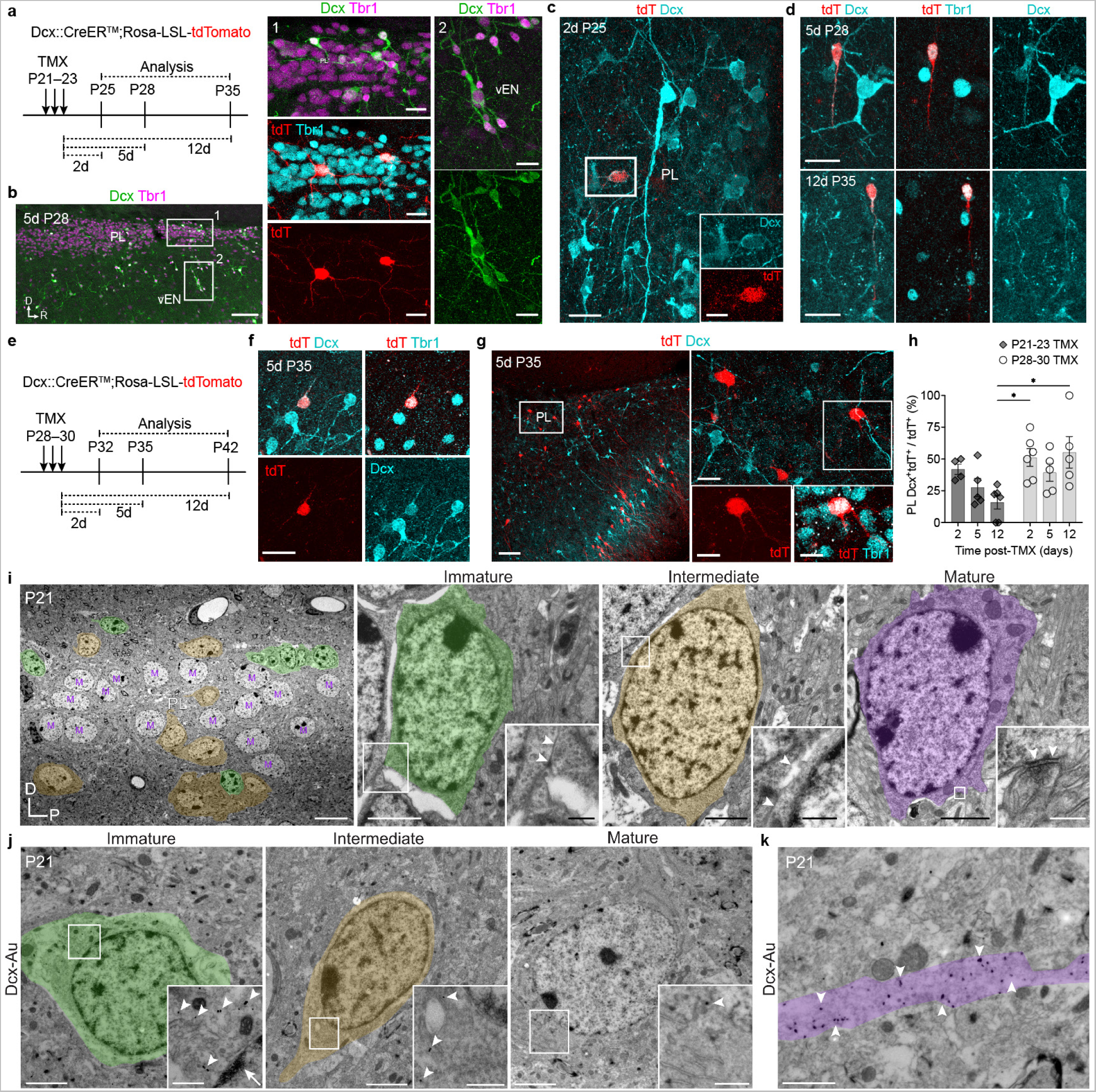
PL neurons downregulate Dcx expression as they mature. **a,** Schematic of experimental design for early labeling of Dcx-CreER^TM^;Rosa-LSL-tdTomato mice. TMX was given once per-day on P21, P22, and P23 and brains were examined either 2, 5, or 12 days following the final dose. **b,** Examples of Dcx^+^ cells with varying morphology in the PL and vEN at P28 (5-day chase). Two tdT^+^Tbr1^+^ cells are present in the PL with large somas and complex arborization and are Dcx–. **c,** Example of Dcx^+^tdT^+^ cell in the PL with a small soma and few branches at P25 (2-day chase). **d,** Examples of Dcx^+^tdT^+^Tbr1^+^ cells in the PL at P28 (5-day chase) and at P35 (12-day chase) with a small elongated soma and one extended process with no visible branching. **e,** Schematic of experimental design for late labeling of Dcx-CreER;ROSA-LSL-tdTomato mice. TMX was given once per-day on P28, P29, and P30 and brains were examined either 2, 5, or 12 days following the final dose. **f,** Example of Dcx^+^tdT^+^Tbr1^+^ cell in the PL with a small soma and few branches at P35 (5-day chase). **g,** Examples of Dcx^+^ cells with varying morphology in the PL and vEN at P35 (5-day chase). Two tdT^+^Tbr1^+^ cells are present in the PL with large somas and complex arborization and are Dcx^−^. **h,** Quantification of the percentage of tdT^+^ cells in the PL that co-express Dcx at each chase timepoint following early TMX (diamonds) or late TMX (circles) injections. **i,** Transmission electron microscopy of the PL nucleus at P21 showing a high density of mature neurons (M) surrounded by intermediate (yellow) or immature neurons (green). Immature neurons have intercellular space and zonula adherens-like contacts (arrows). Mature PL neurons have axosomatic synapses (arrowheads) with pleomorphic vesicles. **j,** Dcx-immunogold (arrowheads) labeled PL neurons. Immature neurons (green) have compacted chromatin (arrow), limited cytosol and organelles (inset). Mature neurons (purple) have more organelles and less heterochromatin and Dcx-labeling. **k,** Dcx-immunogold (arrowheads) in a cell expansion (pseudocolored green). * = p < 0.05, 2-way ANOVA with Holm-Sidak’s post-hoc test. Scale bars = 100 µm (**b** left, **g** left), 20 µm (**b** middle and right, **c** top, **d, f, g** right), 10 µm (**c** bottom, **i** left overview), 2 µm (**i** right overviews, **j** overviews), 1 µm (**j** mature inset, **k**), 500 nm (**i** and **j** immature and intermediate insets), 200 nm (**i** mature inset). PL: paralaminar nucleus; vEn: ventral endopiriform cortex; tdT: tdTomato; TMX: tamoxifen; M: mature neuron.

To further investigate local maturation of PL neurons, we used transmission electron microscopy to study their ultrastructure (**Fig. 4i–k**). At P21, the PL is composed of a dense collection of mature neurons intermingled with neurons of smaller size and with immature ultrastructural features (**Fig. 4i,j**). Immature neurons had compacted chromatin and limited cytosol and few organelles other than ribosomes. Interestingly, some of these cells had intercellular space and zonula adherens-like contacts with neighboring cells, and Dcx-immunogold labeled expansions (**Fig. 4k**). Some neurons had more cytosol and organelles, but retained some compacted chromatin, appearing to be in intermediate stages of maturation. The most mature PL neurons exhibited more mitochondria and larger endoplasmic reticulum, and axosomatic synapses containing pleomorphic vesicles. Together these findings support the conclusion that the PL contains a heterogeneous collection of immature neurons in different stages of maturation, including a subpopulation of immature neurons that downregulate Dcx as they mature and a subpopulation that retains Dcx expression.

### Tbr1^+^ neurons with migratory morphology emanate from the PL during adolescence

In our initial investigations we observed a field of Dcx^+^ cells extending from the PL into the vEN at P21 (**Figs. 1a,b, 2a, 3a**). At this same age, total Dcx^+^ cells dropped sharply in the PL alongside an increase in the vEN (**Fig. 3**), suggesting the possibility of an adolescent wave of neuron migration from the PL to the vEN. To investigate this, we first examined the morphology of these Dcx^+^ cells. To distinguish them from the migratory interneurons in the RMS which express Sp8^56^, we co-stained for this marker alongside Tbr1 which is expressed by a large fraction of PL Dcx^+^ cells (**Fig. 2b**). We found a large population of Dcx^+^ neurons with migratory features and nearly all of these cells were Dcx^+^Tbr1^+^, with few Dcx^+^Sp8^+^ cells except in the most medial sections near the RMS (**Fig. 5, Extended Data Fig. 5, 6a,b**). We also observed similar small, elongated tdT^+^Tbr1^+^Dcx^+^ cells in the Dcx-CreER^TM^ reporter mice given TMX at P21–P23 or at P28–P30 (**Fig. 5b, Extended Data Fig. 4c,d**) and in Dcx-mRFP mice (**Extended Data Fig. 6c**). The ultrastructure of immature PL neurons at P21 revealed morphologies consistent with migration, including fusiform shapes, microtubule-filled leading processes, spaces between cells, small simple organelles, and limited cytosol (**Fig. 5c,d, Extended Data Fig. 6d,e**).

**Figure 5:**
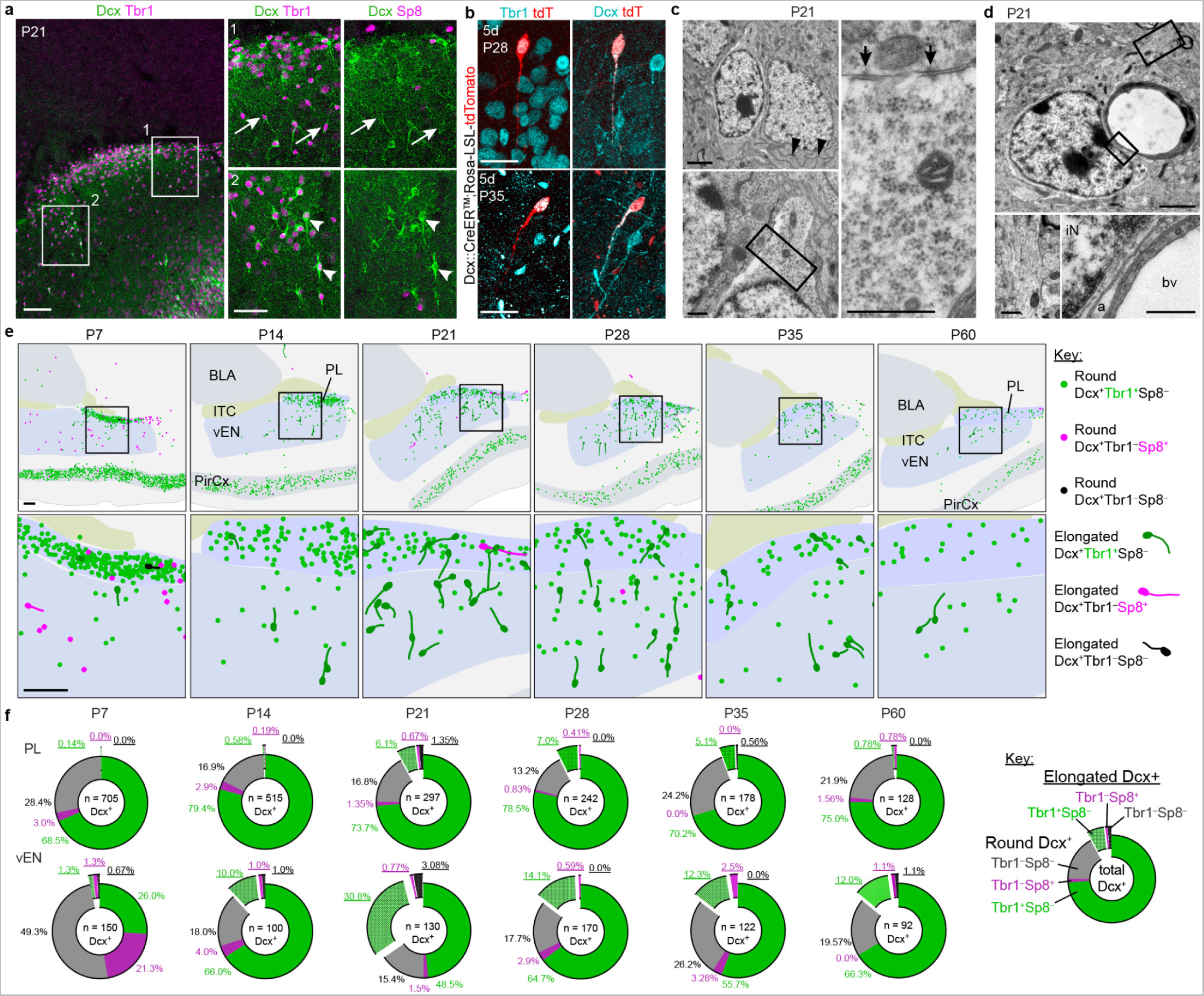
A wave of Tbr1^+^ PL neurons with migratory features appears in adolescence. **a,** Immunostaining of Dcx^+^ cells at P21 reveals a subset that are more rounded/complex Dcx^+^Tbr1^+^Sp8^−^ (arrowheads) and a subset that are more elongated (arrows). **b,** Elongated Dcx^+^Tbr1^+^ cells were labeled with tdT in Dcx-CreER;ROSA-LSL-tdTomato reporter mice at both early (P21-23) or late (P28-30) timepoints. **c,** Ultrastructure of a fusiform neuron in the PL at P21 showing characteristic features of a migratory immature neuron including compacted chromatin, an elongated nucleus and cell body compared to an adjacent more mature neuron with long endoplasmic reticulum (black arrowheads). This cell has few organelles aside from ribosomes (inset) and cell-cell contacts typical of a migratory neuron (arrows) and intercellular spaces. **d,** Ultrastructural details of an immature neuron (iN) associated with a blood vessel (bv) in the PL at P21. This cell has an expansion with microtubules (inset) and a close association with an astrocytic process (a) next to the blood vessel. **e,** Maps of the location of round and elongated Dcx^+^ cells co-expressing either Tbr1 (green) or Sp8 (magenta) or neither (black) between P7 and P60 in a 1.5 mm square. The morphologies of fusiform Dcx^+^ cells were traced, indicating the orientation of the elongated process. **f,** Quantification of the percentage of round and elongated Dcx^+^ cells co-expressing either Tbr1 (green) or Sp8 (magenta) or neither (black) between P7 and P60 in the PL (top row) and vEN (bottom row). Elongated percentages are underlined and popped out of the chart. Total number of Dcx^+^ cells evaluated in each region and age is indicated in the center of each chart. Scale bars = 100 µm (**a** left, **e**), 50 µm (**a** right insets), 20 µm (**b**), 2 µm (**c** top left, **d** top), 500 nm (**c** bottom and right, **d** bottom). PL: paralaminar nucleus; vEN: ventral endopiriform cortex; BLA: basolateral amygdala; iN: immature neuron; a: astrocyte; bv: blood vessel.

Next, we mapped the location and shape of all Dcx^+^ neurons with elongated profiles from P7-P60 within a 1.5 mm square centered on the PL using Neurolucida (**Fig. 5e**, additional levels in **Extended Data Fig. 6f**). We observed few elongated Dcx^+^Tbr1^+^ cells in the PL and vEN at P7 or P14, and a sharp increase in their presence between P21 and P28 (**Fig. 5e, Extended Data Fig. 6f,h–k**). At this peak, the percentage of all Dcx+ cells that were Dcx^+^Tbr1^+^ elongated neurons was as high as 6-7% in the PL and 30% in the vEN (**Fig. 5f**); in contrast, few Sp8^+^Dcx^+^ cells were observed in either region at any age (less than 1%, **Fig. 5f**). At P21 and P28, the leading processes of the elongated Dcx^+^Tbr1^+^ cells were mostly oriented radially between the vEN and the PL (**Fig. 5e**). Fewer elongated Dcx^+^Tbr1^+^ cells were observed at P35 and P60 (**Fig. 5f**). We also detected very few cells with elongated morphology near the piriform cortex or BLA, or entering the PL from other nearby areas in the sections we mapped (**Fig. 5e, Extended Data Fig. 6f,g,j,k**). Together with the timing of cell population changes (**Fig. 3**), these observations support the conclusion that a subpopulation of Dcx^+^Tbr1^+^ neurons migrate from the PL into the vEN postnatally. Based on this surprising presence of a population of immature migratory neurons postnatally, we next asked if, like the RMS, any of these cells were produced via postnatal neurogenesis.

### Postnatal birth and death of neurons in the PL is rare

The presence of immature neurons in the mouse subgranular zone^57,58^ and subventricular zone^59^ is a consequence of postnatal neurogenesis. To determine when immature Dcx^+^ neurons in the PL or vEN are born, we conducted BrdU labeling. We administered BrdU once-daily between P10–P16, a time period prior to the decline of PL Dcx^+^ cells and migration into the vEN. We quantified the density of BrdU-labeled cells co-stained for Dcx or NeuN in the PL and vEN after a 5-day delay (at P21) or 19-day delay (at P35) to permit time for cellular maturation (**Fig. 6**). We found no evidence of BrdU-labeled NeuN^+^ cells at either age (**Fig. 6a–e,i–k**). Some BrdU signal initially appeared to overlap with NeuN, but orthogonal views of the z-stacks revealed that the BrdU was non-overlapping (**Fig. 6j,k**), possibly representing dividing satellite glial cells^44^. Interestingly, we observed a low frequency (~1.3%) of BrdU^+^ cells that were Dcx^+^NeuN^−^ in the PL (**Fig. 6h,i,k**); the percentage and density of these cells was constant in the PL and vEN and at either age examined (**Fig. 6f,g**). The few BrdU^+^ cells that were Dcx^+^NeuN^−^ did not have the elongated morphologies of migratory cells (**Fig. 6i,f,g**), and instead were small and rounded.

**Figure 6:**
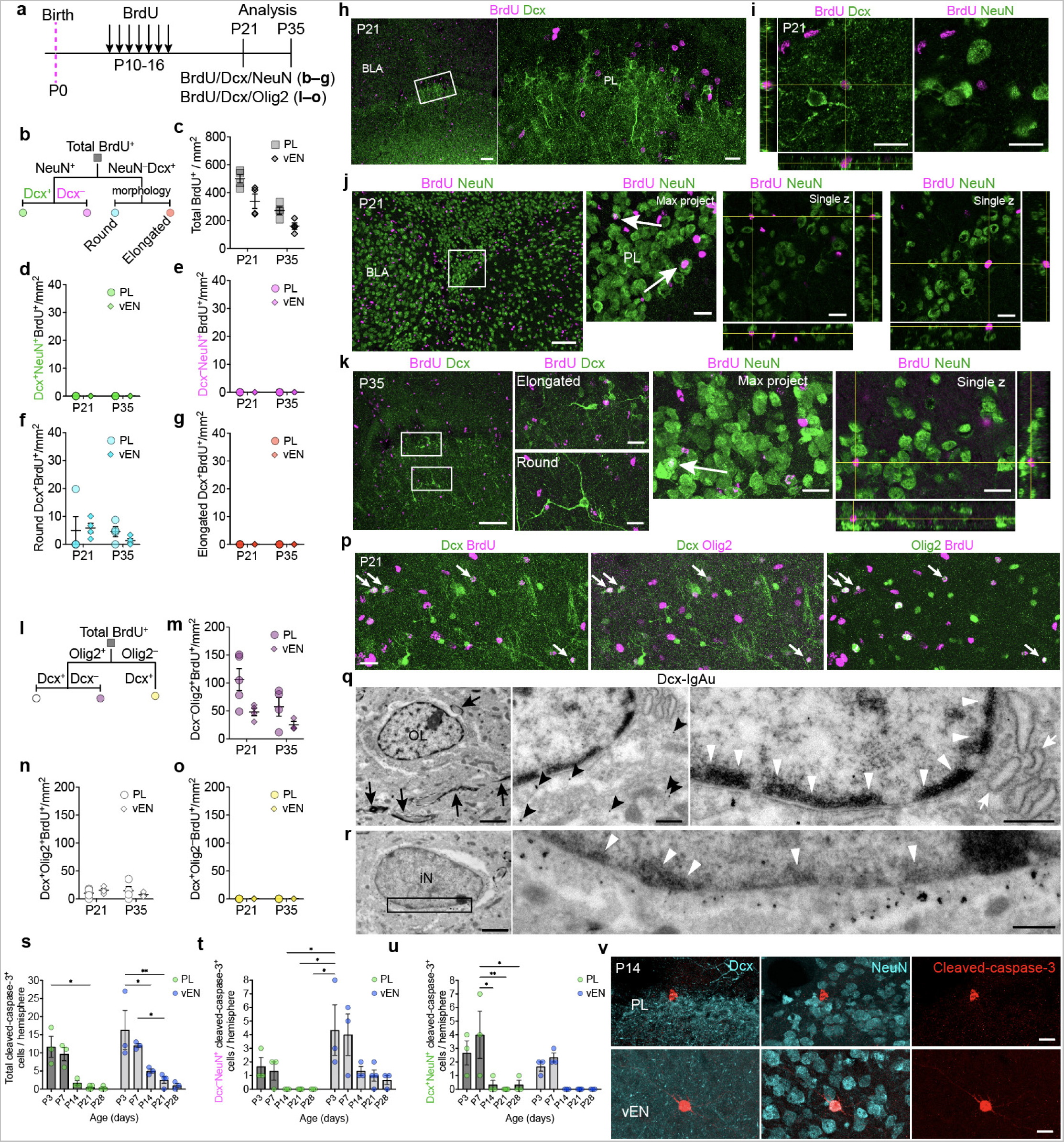
Neuron birth and death are rare in the postnatal PL and vEN. **a,** BrdU was given once per-day from P10 to P16 and brains were examined at P21 or P35. **b,** Cell types analyzed from sections co-stained for BrdU, Dcx, and NeuN. The NeuN^+^ fraction was split into Dcx^+^ cells and Dcx^−^ cells. In a separate analysis the Dcx^+^ cell population was separated by morphology into round and elongated subtypes. **c–g,** The density of all BrdU^+^ cells (**c**), Dcx^+^NeuN^+^BrdU^+^ cells (**d**), and Dcx^−^NeuN^+^BrdU^+^ cells (**e**) was quantified at P21 and P35 in the PL and vEN. Density of Dcx^+^BrdU^+^ cells with round (**f**) or elongated morphology (**g**). **h,** Representative immunostaining of BrdU and Dcx at P21. **i,** Example of round Dcx^+^NeuN^−^ cell co-expressing BrdU in the PL. **j,** Representative immunostaining of BrdU and NeuN in the PL at P21. Putative double-positive cells (arrows) are determined with orthogonal views of the z-stack to be single-positive despite the close proximity of BrdU and NeuN immunostaining signal. All putative double-positive cells for quantifications (**d–g, m–o**) were confirmed this way. **k,** Representative immunostaining for BrdU, Dcx, and NeuN at P35 in the PL and vEN. Examples of BrdU^−^ elongated and round Dcx^+^ cells. In the top panel, one BrdU^+^ cell overlapped with NeuN (arrow) in the maximum intensity projection of the z-stack and was determined via orthogonal views to be single-positive. **l,** Cell types analyzed from sections co-stained for BrdU, Dcx, and Olig2. The Olig2^+^ subset was split into Dcx^+^ cells and Dcx^−^ cells and considered alongside the Olig2^−^Dcx^+^ fraction. **m–o,** The density of Dcx^−^Olig2^+^BrdU^+^ cells (**m**), Dcx^+^Olig2^+^BrdU^+^ cells (**n**), and Dcx^+^Olig2^−^BrdU^+^ cells (**o**) was quantified at P21 and P35 in the PL and vEN. **p,** Example of four BrdU^+^Olig2^+^Dcx^+^ cells (arrows) in the PL at P21. **q,** Ultrastructural details of a weakly Dcx immunogold (black arrowheads) labeled cell displaying typical features of an oligodendrocyte (OL) including nearby myelin (black arrows), dilated, short endoplasmic reticulum (white arrows), and distinctive condensed chromatin along the nuclear envelope (white arrowheads). **r,** Ultrastructural details of a strongly Dcx immunogold labeled cell displaying typical features of an immature neuron (iN) including a fusiform shape, limited cytosol, few organelles, and condensed chromatin (white arrowheads). **s–u**, Quantifications of the total number of cleaved-caspase-3^+^ (cc3^+^) (**s**), Dcx^−^NeuN^+^cc3^+^ (**t**), and Dcx^+^NeuN^+^cc3^+^ cells (**u**) in the PL (green) or vEN (blue) between P3 and P28. **v,** Example immunostaining of cleaved-caspase-3^+^ cells in the PL (Dcx^−^ and NeuN^−^) and vEN (Dcx^−^, NeuN^+^). * = p < 0.05, ** = p < 0.01, *** = p < 0.001. 2-way ANOVA with Holm-Sidak’s post-hoc test. Scale bars = 100 µm (**h** left, **j** left, **k** left), 20 µm (**h** right, **i, j** right, **k** right, **p, v**), 2 µm (**q** left, **r** left), 500 nm (**q** middle and right, **r** right). BLA: basolateral amygdala; PL: paralaminar nucleus; vEN: ventral endopiriform cortex; OL: oligodendrocyte; iN: immature neuron.

To further investigate this small population of Dcx^+^BrdU^+^ cells that was NeuN^−^, we next tested the possibility that these cells might be glia. Dcx can be expressed by OPCs^39–41,43,60^, so we stained PL sections for Dcx alongside the oligodendrocyte markers Olig2 and adenomatous polyposis coli (APC). A subset of Dcx^+^ cells co-localized with Olig2 and APC, and there was little Dcx co-localization with markers of microglia (Iba1) or astrocytes (GFAP) (**Extended Data Fig. 7a–n**). Next, we co-stained the Dcx^+^BrdU^+^ cells in the mice given BrdU between P10-P16 for Olig2. At both P21 and P35 in both the PL and vEN, we frequently observed BrdU^+^Olig2^+^ cells, and a subpopulation were also Dcx^+^. In these brains we did not detect any BrdU^+^Dcx^+^ cells that were Olig2^−^ (**Fig. 6l–p**). At P21 we could readily identify Dcx-immunogold labeled cells in the PL with ultrastructural features distinctive of both oligodendrocytes and immature neurons (**Fig. 6q,r**). Together these results provide evidence for ongoing postnatal gliogenesis, but an absence of postnatal neurogenesis in the PL or vEN.

In the cerebral cortex, there is a peak of developmental programmed cell death that occurs during the first postnatal week^61,62^. Because the developmental timeline of PL neurons is delayed, we predicted that their developmental programmed cell death might also be delayed, possibly occurring during the peak decline of Dcx^+^ cells in the region (P21–P28). To investigate this possibility, we examined every section containing the PL or vEN from mice between P3–P28, stained for Dcx, NeuN, and cleaved-caspase-3 (CC3) to label cells undergoing apoptosis^63^. The peak CC3 expression in the PL and vEN occurred at ages typical for cortical neurons (P3–P7), and was similar for Dcx^−^NeuN^+^ mature neurons, or Dcx^+^NeuN^+^ immature neurons (**Fig. 6s–v, Extended Data Fig. 7o,p**). At all older ages (P14-P28) we observed at most 1-2 CC3^+^ cells per-hemisphere in the PL or vEN. From these observations we conclude that the majority of the changes in numbers of Dcx^+^ and NeuN^+^ cells in the PL and vEN are due to a downregulation of Dcx expression with maturation or migration, but not cell birth or death.

### Immature PL neurons across postnatal ages and morphologies share a common embryonic birthdate

As PL neurons are not born postnatally, we next investigated their embryonic birthdate. We administered BrdU to timed-pregnant female mice once between E12.5 – E16.5, or pups at P5. At P21, we quantified the percentage and density of Dcx^+^NeuN^+^ immature neurons co-expressing BrdU and found that their peak birth-date was E14.5 in both the PL and vEN, with no evident postnatal contribution (**Fig. 7a–d, Extended Data Fig. 8a–f**). Interestingly, this peak birthdate was the same for the Dcx^+^NeuN^+^ PL and vEN neurons at P7 and at P35 (**Fig. 7e–h**), revealing that immature neurons at all postnatal ages have the same embryonic birthdate, and therefore retain Dcx expression for weeks into adolescence before maturing or migrating. Furthermore, both PL and vEN immature neurons shared the same embryonic birthdate, indicating that despite their abundance in both regions, neither region is a postnatal birthplace for immature neurons. Consistently, the mature (Dcx^−^NeuN^+^) neurons in the PL were born embryonically and later than the majority of the vEN mature neurons (**Fig. 7i–k**).

**Figure 7:**
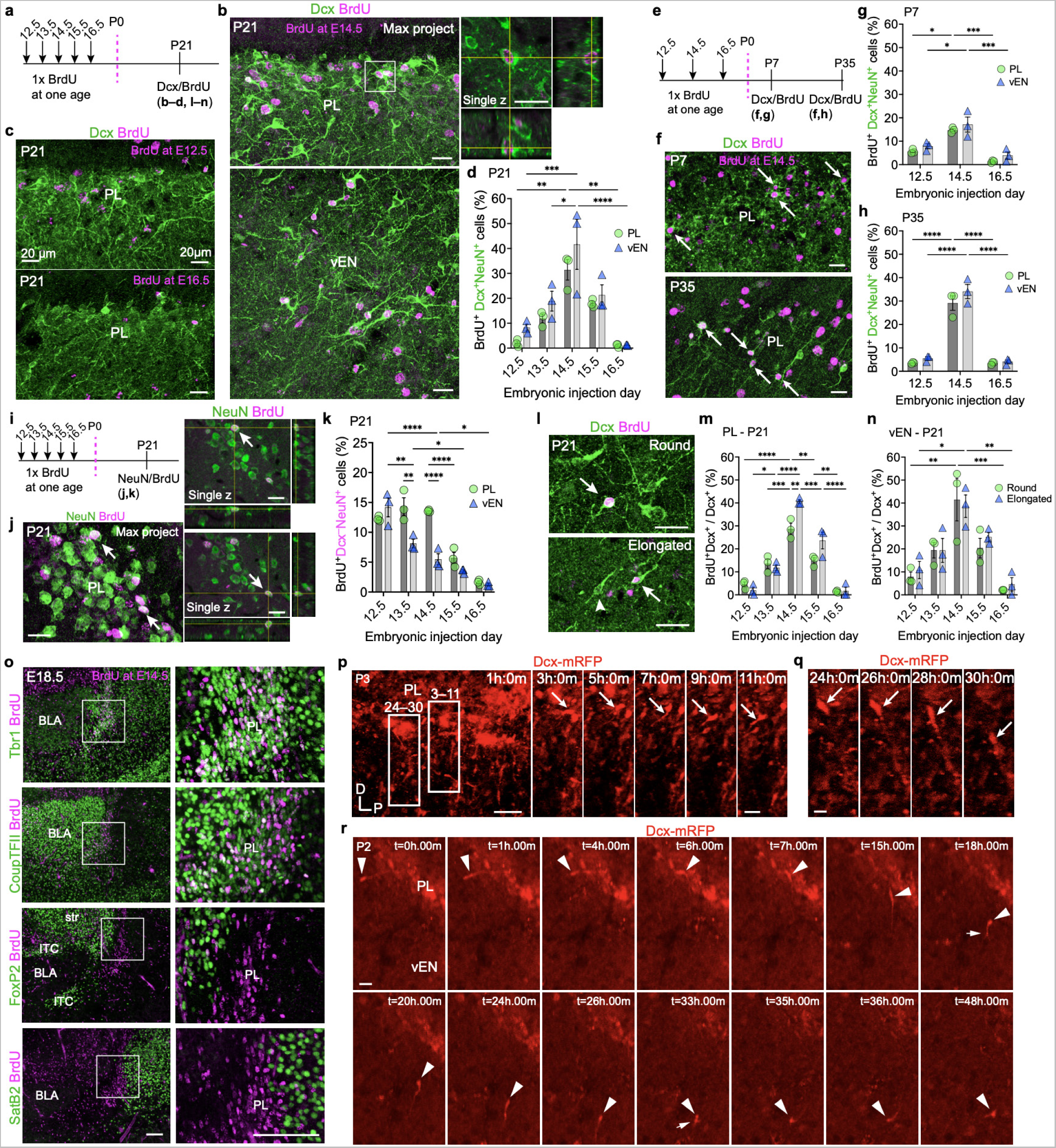
Neurons in the PL are born during embryonic ages. **a,** BrdU was given once to timed-pregnant mice between E12.5 – E16.5 and offspring were examined at P21. **b,** Examples of BrdU^+^Dcx^+^ cells in the PL and vEN at P21. Orthogonal views of the z-stack show that the BrdU^+^ signal is contained within the nucleus of the Dcx^+^ cell. **c,** Infrequent Dcx^+^ cells in the PL at P21 from either E12.5 or E16.5 BrdU. **d,** Quantification of the percentage of Dcx^+^NeuN^+^ cells in the PL and vEN at P21 that are BrdU^+^ from a single injection between E12.5 – E16.5. **e,** BrdU was given once to timed-pregnant females between E12.5 – E16.5 and brains were examined at P7 or P35. **f,** Examples of BrdU^+^Dcx^+^ cells (arrows) in the PL at P7 and P35. **g,h,** Quantification of the percentage of Dcx^+^NeuN^+^ cells in the PL and vEN at P7 (**g**) or P35 (**h**) that are BrdU^+^ from a single injection between E12.5 – E16.5. **i,** BrdU was given once to timed-pregnant mice between E12.5 – E16.5 and offspring were examined at P21. **j,** Examples of BrdU^+^NeuN^+^ cells in the PL at P21 (arrows). Orthogonal views of the z-stack show that the BrdU^+^ signal overlaps with NeuN. **k,** Quantification of the percentage of Dcx^−^NeuN^+^ cells in the PL and vEN at P21 that are BrdU^+^ from a single injection between E12.5 – E16.5. **l,** Examples of BrdU^+^Dcx^+^ cells classified as either round or elongated (arrows) near an elongated BrdU^−^Dcx^+^ cell (arrowhead). **m,** Quantification of the percentage of round (green) or elongated (blue) Dcx^+^ cells that co-express BrdU from indicated injection ages at P21 in the PL (**m**) or vEN (**n**). **o,** At E18.5, following an E14.5 BrdU injection, BrdU^+^Tbr1^+^ cells and BrdU^+^CoupTFII^+^ cells are present in a dense region anterior-ventral to the BLA between a BrdU^−^FoxP2^+^ region and a BrdU^−^SatB2^+^ region. **p–r,** Acute slice preparations from P2-3 Dcx-mRFP mice showing individual frames at indicated timestamps from a time-lapse confocal image acquisition series. **p,r,** The initial timelapse field is shown 1 hr after imaging began. Five frames of hours 3–11 (**p**) show a cell extending its process from the PL into the nearby vEN (arrow). Four frames from hours 24–30 (**r**) show a neuron that was previously stationary within the PL beginning to migrate away from the PL (arrow). **r,** Fourteen frames from a separate acquisition show a Dcx-mRFP^+^ cell migrating into the PL from the nearby cortex (arrowhead). At t = 7.00 hrs, this cell re-orients by 90-degrees and exits the PL. At t = 18.00 hrs, the cell has a bifurcated leading process (arrow) prior to resuming migration deeper into the cortex. This bifurcation occurs again at t = 33.00 hrs prior to the cell appearing to settle and ramify between t = 35.00 hrs and t = 48.00hrs. * = p < 0.05, ** = p < 0.01, *** = p < 0.001, **** = p < 0.0001. 2-way ANOVA with Holm-Sidak’s post-hoc test. Scale bars = 100 µm (**o**), 50 µm (**p** left overview), 20 µm (**b**,**c**,**f**,**i**,**m, p** right, **r**), 10 µm (**q**). PL: paralaminar nucleus; vEN: ventral endopiriform cortex, BLA: basolateral amygdala.

Although they appeared to emerge from the dense collection of round neurons in the PL, the elongated Dcx^+^ cells could be migrating away from a postnatal birthplace similar to known neurogenic regions^28,30^. To investigate this possibility we next examined the birthdate of Dcx^+^ cells with migratory morphology. We found that the peak birthdate for both round and elongated neurons was E14.5, and was similar between the PL and vEN at every postnatal age examined (**Fig. 7l–n, Extended Data Fig. 8g–o**). Thus, despite their particularly immature features, these elongated migrating neurons are born embryonically, indicating that they remain arrested in their development for prolonged periods of time following their genesis before unknown factors stimulate them to migrate during adolescence. These results also point to the possibility of a shared embryonic origin for the immature neurons in both the PL and vEN.

### The PL forms during embryonic ages in mice

Since the PL is visible postnatally as early as P3 (**Fig. 2j**), and its neurons are born embryonically, we next investigated the earliest age that we could identify this region. At E16.5, a region with Tbr1^+^CoupTFII^+^ cells was not present near FoxP2^+^ or SatB2^+^ cells; however at E18.5 a dense nucleus of cells was visible anterior and ventral to the BLA (**Fig. 7o, Extended Data Fig. 9a,b**). Following E14.5 BrdU injection, this region contained BrdU^+^Tbr1^+^ cells and BrdU^+^CoupTFII^+^ cells, but few BrdU^+^SatB2^+^ or BrdU^+^FoxP2^+^ cells, similar to the expression profile of the postnatal PL. Together these data indicate that many of the cells in the mouse PL are established by E18.5. Interestingly, we noticed some BrdU^+^Tbr1^+^ cells emanating ventrally out of the PL at this early age, so we next investigated whether these cells might be migrating using acute slice time-lapse imaging.

We prepared acute slice cultures containing the PL from P2–P3 Dcx-mRFP mice and imaged them up to 48 hrs with a confocal microscope (**Fig. 7p–r, Supplementary Movie 3**). In these sections we observed many examples of migratory mRFP^+^ cells with their cell bodies originally located within the PL, extending their leading process into the neighboring vEN (**Fig. 7p**). We also observed a neuron within the PL that was stationary for the first 24 hrs of the imaging before it initiated migration from the PL into the vEN (**Fig. 7q**). In a separate section we followed an mRFP^+^ cell with its soma initially located in the vEN extending its leading process into the PL, reorienting, and re-entering the vEN where its leading process bifurcated multiple times before choosing a final location to settle after 48 hrs (**Fig. 7r**). These data demonstrate that migration from the PL into the vEN occurs postnatally, and the further intriguing observation that neurons located in the PL can initiate migration after up to 24 hrs of remaining stationary.

### The PL forms during embryonic ages in humans

It is not known when the PL is established in humans; however, it has been described to form in the final gestational months^36^. We previously identified a high density of COUPTFII^+^ cells expressing *TBR1* as early as 22 GW in the region where the PL COUPTFII^+^DCX^+^ cells are observed at birth^7^. Using our observations of a mid-embryonic neurogenesis birthdate of the PL in mice and PL expression of CoupTFII, Ctip2, and Tbr1, but not SatB2 or FoxP2 in the mouse PL, we next explored when this region forms during human gestation.

To investigate the earliest ages for possible formation of the human PL, we prepared coronal and horizontal sections from an 18 GW embryo and stained serial sections for the positive and negative PL markers (**Fig. 8a–c, Extended Data Fig. 10a,b**). In two sections containing the most ventral extension of the temporal lobe lateral ventricle, we identified a thin stripe of COUPTFII^+^TBR1^+^ cells in the anatomical location of the PL at later ages, which also co-labeled with CTIP2 but not SATB2 or FOXP2. We observed the same anatomical expression pattern at 22 GW, with the density of COUPTFII^+^TBR1^+^ cells extending more anteriorly, and growing in size along the wall of the ventricle (**Fig. 8d–f**). At 28 GW, the high density of cells in the PL was distinctive compared to the rest of the BLA, and looked similar to the shape and anatomical distribution of the cells at birth (**Fig. 8g–l, Extended Data Fig. 10c–e**). At birth, these neurons extended rostral to the BLA, similar to what is observed in mice (**Extended Data Fig. 2, 10c**). Between 27 GW and 13 years of age, TBR1, COUPTFII, and CTIP2 labeled the majority of DCX^+^ cells in the PL, whereas few cells were labeled by FOXP2, SATB2, or GAD67 (**Fig. 8m,n**). Interestingly, while we have previously observed DCX^+^ cells in the PL with elongated morphology across human lifespan^7^, it was not known whether these cells were associated with the immature PL neurons. Using the transcription factors we identified as highly expressed in immature PL neurons, we observed similar elongated DCX^+^ cells that were TBR1^+^COUPTFII^+^, or CTIP2^+^SATB2^−^ at 13 years of age (**Fig. 8o,p**). Together these data suggest that the cells comprising the human PL begin to appear as early as 18 GW, extend rostrally with increasing gestational age, and share molecular, morphological, and developmental features with the mouse PL.

**Figure 8:**
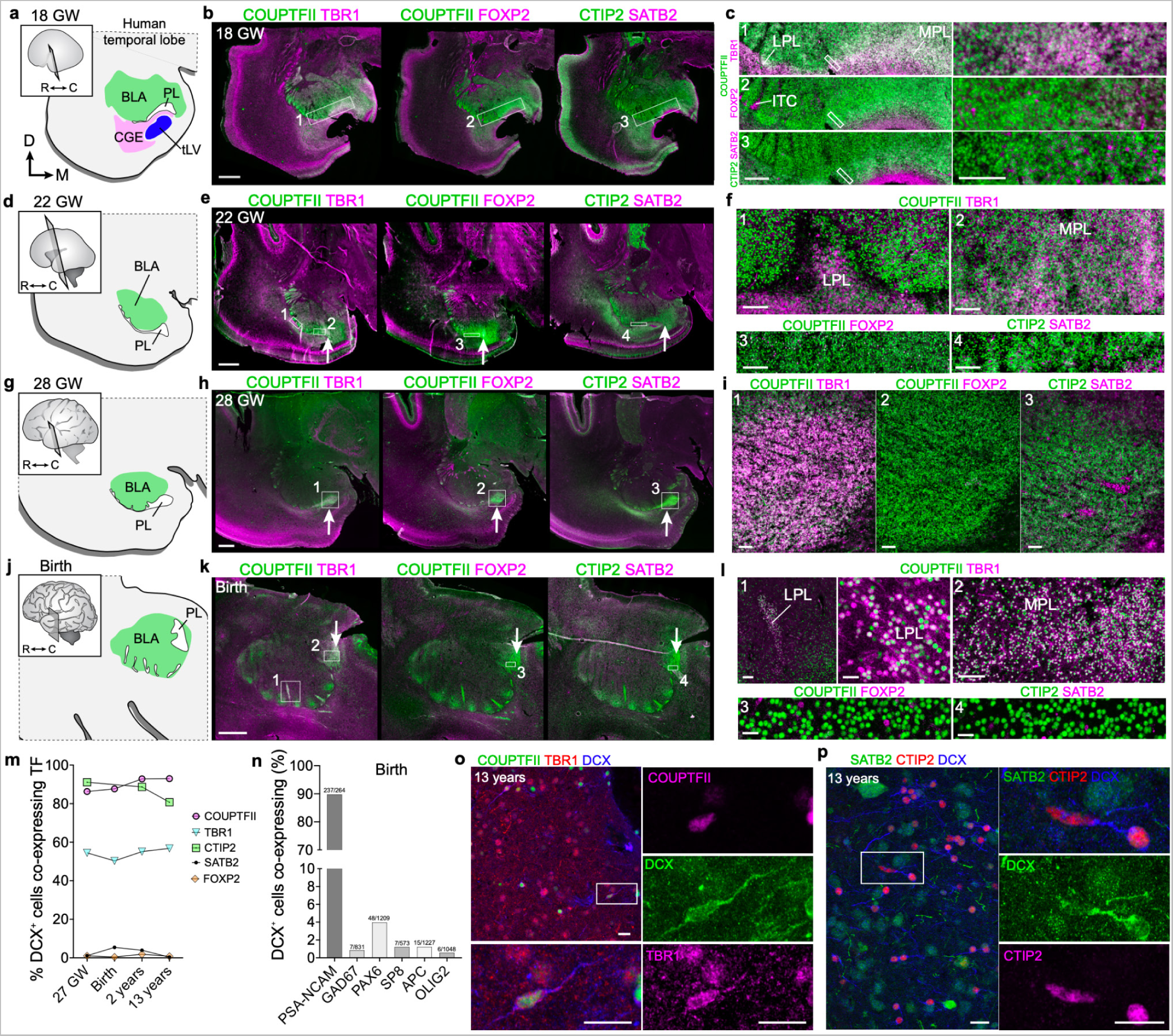
The human PL forms between GW 18 and 22. **a–c,** 18 GW human temporal lobe coronal sections stained for COUPTFII and TBR1 (1), COUPTFII and FOXP2 (2), or CTIP2 and SATB2 (3). Map (**a**) indicating the location of the BLA relative to the PL and temporal lobe lateral ventricle (tLV). Overviews of the immunostaining across each section (**b**) showing the location of the boxed inset regions in (**c**). **d–f,** 22 GW human temporal lobe coronal sections stained for COUPTFII and TBR1 (1,2), COUPTFII and FOXP2 (3), or CTIP2 and SATB2 (4). Map (**d**) indicating the location of the BLA relative to the PL which is now visible at this age at levels immediately rostral to the tLV. Overviews of the immunostaining across each section (**e**) showing the location of the boxed inset regions in (**f**) and the PL (arrows). **g–i,** 28 GW human temporal lobe coronal sections stained for COUPTFII and TBR1 (1), COUPTFII and FOXP2 (2), or CTIP2 and SATB2 (3). Map (**g**) indicating the location of the BLA relative to the PL which is visible at this age at rostral levels. Overviews of the immunostaining across each section (**h**) showing the location of the boxed inset regions in (**i**) and the PL (arrows). **j–l,** Birth human temporal lobe coronal sections stained for COUPTFII and TBR1 (1,2), COUPTFII and FOXP2 (3), or CTIP2 and SATB2 (4). Map (**j**) indicating the location of the BLA relative to the PL. Overviews of the immunostaining across each section (**k**) showing the location of the boxed inset regions in (**l**) and the PL (arrows). **m,** Quantifications of the percentage of DCX^+^ cells in the human PL co-expressing each indicated transcription factor (TF) between 27 GW and 13 years of age. **n,** Quantifications of the percentage of DCX^+^ cells in the PL co-expressing the indicated marker at birth in the human PL. **o,** Example of a small DCX^+^ neuron at 13 years of age in the human PL that has an elongated nucleus, a putative leading process, and co-expresses both COUPTFII and TBR1. **p,** Example of an DCX^+^CTIP2^+^SATB2^−^ neuron with an elongated nucleus at 13 years of age in the human PL that extends its DCX^+^ process intertwined with a neighboring round DCX^+^ cell. Scale bars = 2 mm (**b, e, h, k**), 500 µm (**c** left), 100 µm (**c** right, **f, i, l** top left and top right) 20 µm (**l** top middle, bottom row, **o, p**). GW: gestation week; BLA: basolateral amygdala; PL: paralaminar nucleus; CGE: caudal ganglionic eminence; tLV: temporal lobe lateral ventricle; LPL: lateral paralaminar nucleus: MPL: medial paralaminar nucleus: ITC: intercalated cell cluster; TF: transcription factor.

## Discussion

Immature neurons are present in the postnatal mouse brain in well-described adult neurogenic regions including the subgranular zone^57,58^ and subventricular zone^59^, as well as in adult non-neurogenic regions like the piriform cortex^15,64–66^. Here we show that the adolescent mouse PL is also a non-neurogenic region containing immature Dcx^+^ neurons adjacent to the BLA. Similar to the human PL, these Dcx^+^ cells are immature excitatory neurons that undergo a delayed maturation during early adolescence (P21-28). Their decrease in number was not associated with programmed cell death, but rather with their maturation and corresponding loss of Dcx expression. Strikingly, a subpopulation of the immature excitatory neurons are not stationary, possessing morphological and ultrastructural characteristics of migratory neurons. In a postnatal wave of migration peaking between P21 and P28, these neurons extended their leading process into the neighboring vEN, apparently contributing to an increase in the total number of neurons in the endopiriform cortex. PL neurons were not produced from postnatal neurogenesis; instead, they were born between E12.5 and E14.5, with a peak of E14.5 for those that remained immature into mid-adolescent ages. The transcription factors Tbr1 and CoupTFII distinguished PL neurons from neighboring FoxP2^+^ cells in the ITCs and SatB2^+^ cells in the vEN. This provided a means to identify the earliest ages we could observe PL neurons assemble, when other markers like Dcx are not selective. In mice, we found a dense collection of PL neurons as early as E18.5, and in humans as early as 18 GW. Together these results identify a region adjacent to the mouse BLA that contains neurons with molecular, morphological, and developmental similarity to the human PL. Interestingly, the embryonic birth and formation of the PL suggests that these immature neurons must be either arrested or stimulated by unknown cue(s) that alter the timing of their developmental progression.

The conservation of a collection of immature neurons adjacent to the basolateral amygdala from mice to humans suggests a shared fundamental mechanism of the maturation of the circuits that recruit them during adolescence. In contrast, other regions with delayed-maturation in rodents appear to develop much later in adulthood^64^. In lissencephalic species like bats^67^ and the tree shrew^27^, and mice (**Extended Data Fig. 1d,e, 2**), immature PL neurons are located ventral and lateral to the amygdala, whereas it is present ventral-medial to the BLA in gyrencephalic animals like cats, sheep, and primates^11,18,23^. We observed a rostral extension of the immature PL neurons in mice (**Fig. 2k**) that is anatomically similar to the rostral extension of the larger, possibly more analogous medial PL (MPL) in humans (**Fig. 8, Extended Data Fig 10c,d**). At more caudal levels, the human PL is close to the temporal lobe lateral ventricle (tLV), but in mice the ventricle does not extend beneath the BLA and we did not observe a close association of the PL with the ventricle. The location of the vEN is debated in primates^68^, however some atlases place it between the PL and piriform cortex, a region where, consistent with our observations in mice, a broad field of immature neurons emanate from the PL in humans and non-human primates^7,69^.

The observation of a subset of neurons that display migratory morphology in the mouse PL is consistent with reports of putative migratory cells in the PL of primates^6,7,11,18^, and our molecular identification of cells with this morphology in the 13 year old human (**Fig. 8o,p**). PL neurons are present in a region described as a migratory reservoir in gestational rats^70^ at the ventral extent of the lateral cortical migratory stream, which follows the external capsule^71,72^. Thus, migratory PL excitatory neurons could simply be a continuation of radial migration from embryonic ages. Alternatively, the increase in migration between P21–P28 may be a resurgent second wave of local migration, which is supported by our observation of a neuron in the PL that remained stationary for 24 hours before migrating toward the PL (**Fig. 7p,q**).

The presence of neurons expressing Dcx is often interpreted as evidence of newborn neurons^73,74^, however, our results collectively collectively support an embryonic birthdate for the Dcx^+^ cells in the PL and vEN (**Fig. 7**), with no indications of postnatal neurogenesis. Together our observations indicate that the mouse PL forms during embryonic ages and contains a pool of developmentally-arrested neurons that mature at later ages. Interestingly, a similar process of delayed neuron maturation has been proposed^75^ as an explanation for the presence of DCX^+^ cells with mature morphologies^76^ in the human hippocampus, in the absence of dividing progenitor cells^73,77,78^. A better understanding of the molecular influences on immature neuron maturation arrest and growth might provide a unique angle to understanding the significance of this phenomenon in either brain region.

A powerful type of structural plasticity is the introduction of new neurons into developing or established networks^13^. This “whole-cell” structural plasticity can occur through adult neurogenesis, which in the rodent hippocampus supports processes such as cognitive flexibility and learning^79^. However, most brain regions do not continue to produce newly-born neurons past early postnatal life, especially in mammalian species with large brains^14,73, 80–83^. Regardless of whether an immature neuron is newly born, the process of its growth, maturation, and integration into established circuits is a form of structural plasticity, which is a fundamental mechanism of brain circuit development and adaptation throughout life. In the amygdala these cells develop during critical ages for amygdala-dependent adolescent socioemotional development^6,8,84^. Almost half of all lifetime mental health conditions onset during mid-teenage years^85^, and changes in amygdala volume and development are linked to social and emotional disorders^6,86–90^. Connections between the unique trajectory of PL maturation and neuropsychiatric disorders are beginning to be explored, with some evidence for reduced maturation in MDD^46^ and ASD^6^.

The identification of the PL in mice provides a new focus for investigations into the cellular and molecular development of immature amygdala neurons and the PL as a distinct region compared to the anatomically similar ITCs. Non-newly born immature neurons that progressively grow during critical stages of brain plasticity and development may reveal striking differences with normal neuron functions, or unique traits compared to other regions where these cells are observed.

## Acknowledgments

S.F.S., P.J.A., C.E.P., M.S., D.S., and J.G.C. were supported by the National Institute of Mental Health (NIMH) (R21MH125367 and R01MH128745). S.F.S. and S.W.B. were supported by a Competitive Medical Research Fund award from the University of Pittsburgh Medical Center and start-up funds from the University of Pittsburgh. J.M.G.V. was supported by the Valencian Council for Innovation, Universities, Science and Digital Society (PROMETEO/2019/075) and V.H.P. was supported by the Spanish Ministry of Science, Innovation and Universities (PCI2018-093062). We thank Drs. Julia Kofler and Clayton Wiley, University of Pittsburgh and Eric Huang, University of California, San Francisco for help with human sample collection and the Duke Neurotransgenic Core Facility and Ute Hochgeschwender for assistance in generating the Dcx-CreER^TM^ line. We would also like to thank members of the Corbin lab for constructive feedback through the course of the project and Dr. Arturo Alvarez-Buylla and Quetzal Flores-Ramirez for early discussions and assistance collecting pilot data.

## Author contributions

P.J.A. and S.F.S. conceived the experiments, conceptualized the manuscript and wrote the original draft. P.J.A. performed the mouse immunostainings and confocal microscopy, sterology, BrdU birthdating, time-lapse imaging, and cell mapping under the supervision of S.F.S. Lightsheet imaging and Tshz1-GFP mouse analysis were conducted by D.S. under the supervision of J.G.C. Transmission electron microscopy and analysis were performed by L.T.S. and V.H.P. under the supervision of J.M.G.V. Human tissue samples were processed, stained, and analyzed by M.S., S.W.B. and S.F.S. Mouse embryology experiments were performed by P.J.A. and M.S. and co-localization quantifications in mice were performed by P.J.A., C.E.P., and A.M. under the supervision of S.F.S. Dcx-CreER^TM^ mice and preliminary characterization results were contributed by C.K. Manuscript draft revisions were provided by D.S., C.E.P., V.H.P, and J.G.C. Funding for the study was obtained by J.G.C. and S.F.S. All authors approved the final manuscript.

---

**Extended Data Fig. 1:**
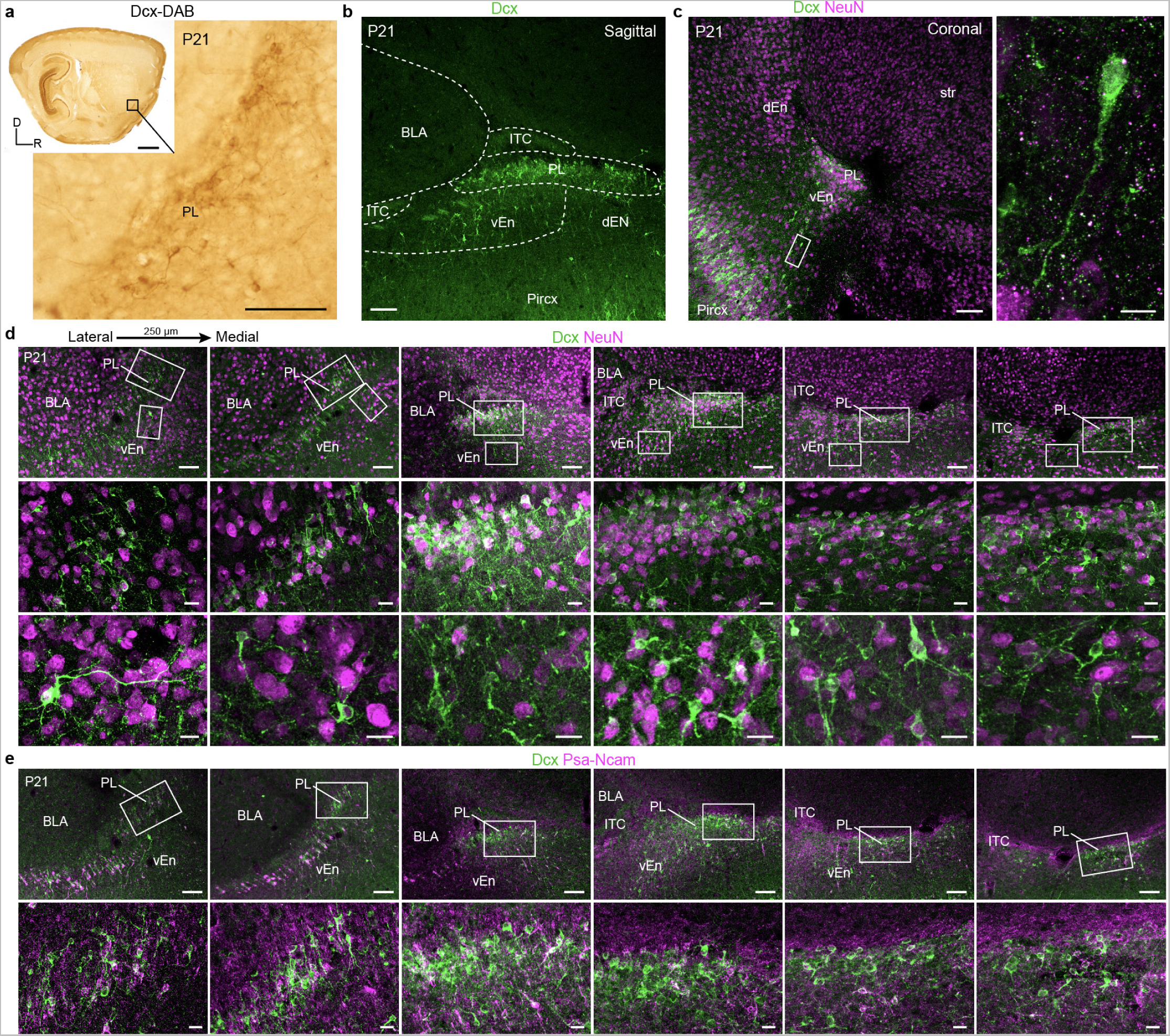
Anatomical characterization of immature neurons in the P21 mouse peri-amygdalar region. **a,** DAB immunohistochemistry of the Dcx^+^ cells in the mouse PL at P21. **b,** Dcx immunofluorescent staining for Dcx in the sagittal P21 section shown in Figure 1a with dotted lines outlining the PL and adjoining regions. **c,** Coronal section of the PL at P21 showing Dcx^+^NeuN^+^ cells and the location of the dEN and vEN. Inset shows an elongated Dcx^+^ cell between the vEN and piriform cortex. **d,** Lateral to medial series of sagittal sections at P21 spaced by 250 µm across the PL. Higher magnification of the PL nucleus and vEN at each level. **e,** Dcx^+^Psa-Ncam^+^ cells within the same section series as in (**d**). Scale bars = 1 mm (**a** top left), 100 µm (**a** bottom right, **b, c** left, **d** top row, **e** top row), 20 µm (**d** middle and bottom rows, **e** bottom row), 10 µm (**c** right). BLA: basolateral amygdala; ITC: intercalated cell clusters; PL: paralaminar nucleus; vEN: ventral endopiriform cortex; dEN: dorsal endopiriform cortex; str: striatum; Pircx: piriform cortex.

**Extended Data Fig. 2:**
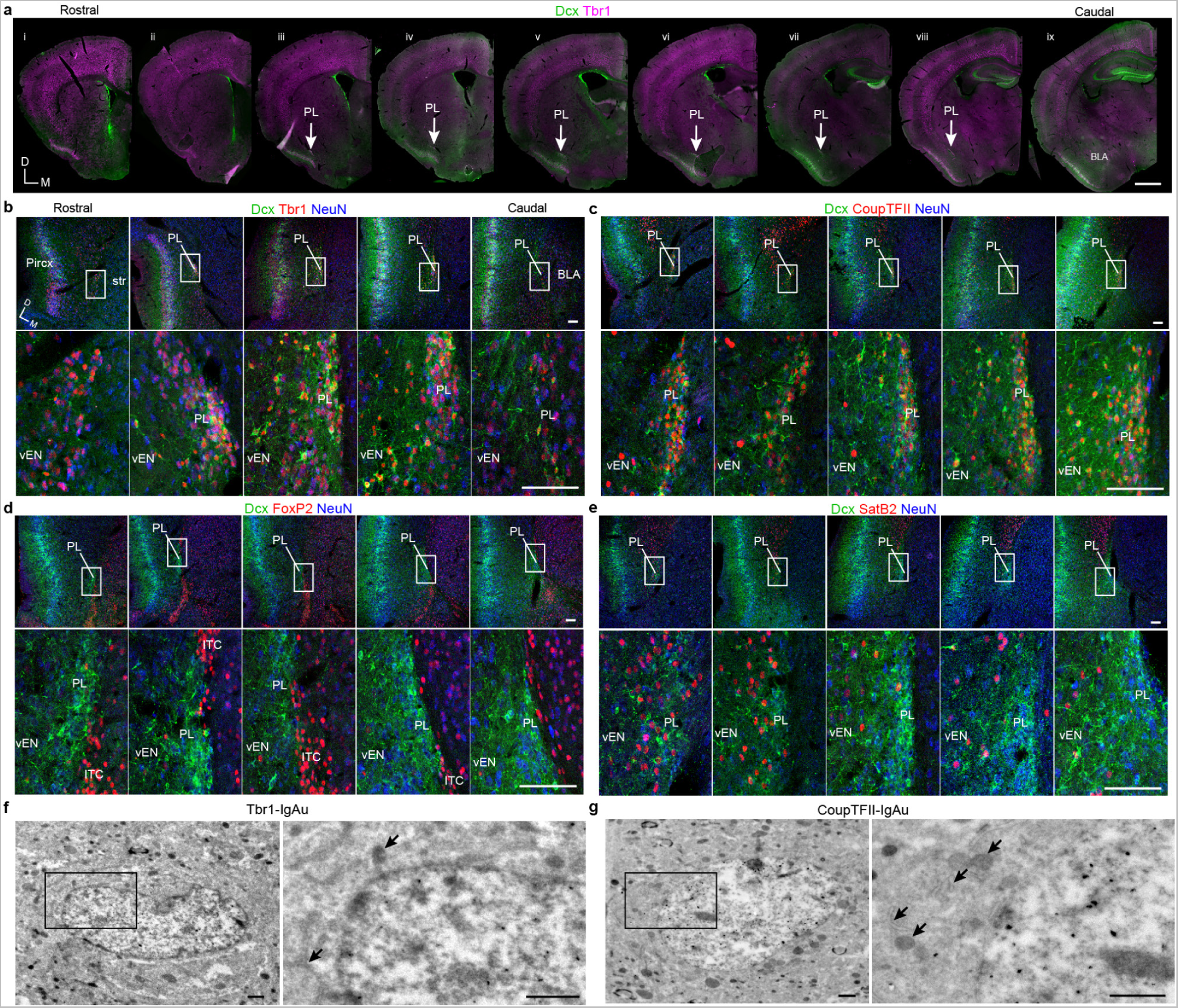
Coronal cross-sections at P21 showing the location of immature neurons in the mouse PL. **a,** Coronal section overviews immunostained for Dcx and Tbr1 showing the location of the PL (arrow) in a rostral-caudal series (spaced by 250 µm). **b–e,** Coronal sections at P21 from 5 rostral-caudal levels with the most caudal section including the BLA. Adjacent sections are stained for Dcx, NeuN, and Tbr1 (**b**), CoupTFII (**c**), FoxP2 (**d**), or SatB2 (**e**). **f,g,** Ultrastructural features of P21 cell immunogold labeled for Tbr1 (**f**) or CoupTFII (**g**). These cells are larger in size and contain more organelles (arrows) and cytosol than immature neurons; the CoupTFII^+^ cell has a slightly elongated profile. Scale bars = 1 mm (**a**), 100 µm (**b–e**), 1 µm (**f**,**g**). Pircx: piriform cortex, str: striatum; BLA: basolateral amygdala.

**Extended Data Fig. 3:**
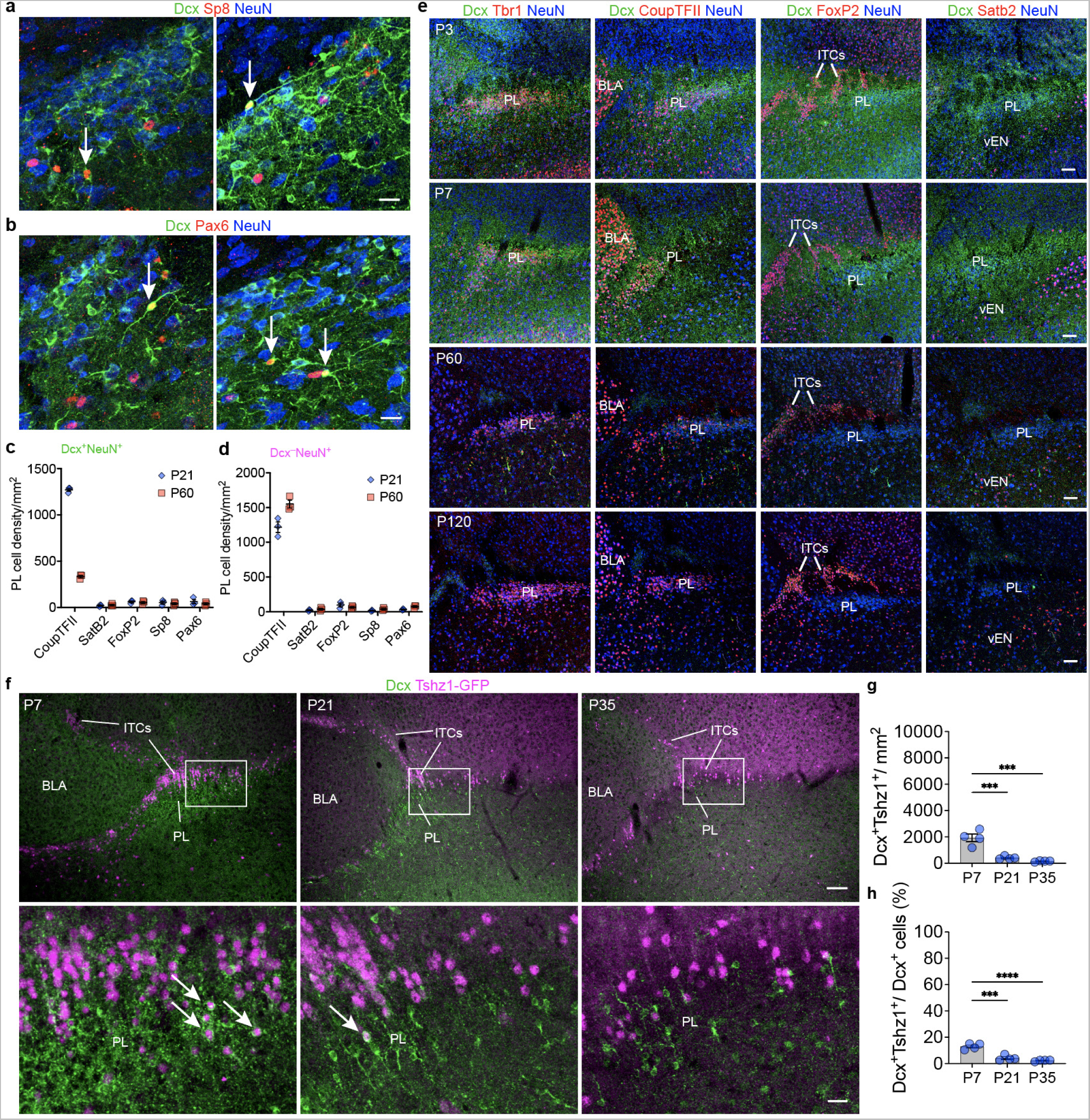
The PL is distinct from the ITCs across postnatal ages. **a,b,** The PL contains small subpopulations of Dcx^+^Sp8^+^ cells (**a**), and Dcx^+^Pax6^+^ cells (**b**) at P21 (arrows). **c,d,** Quantification of the density of Dcx^+^NeuN^+^ cells (**c**) or Dcx^−^NeuN^+^ cells (**d**) co-expressing each indicated transcription factor at P21 or P60. **e,** Age-series of the PL at P3, P7, P60, and P120. At each age, Tbr1^+^ and CoupTFII^+^ cells are present in the PL, FoxP2 cells are present in the ITCs, and SatB2^+^ cells are present in the vEN. **f,** Representative immunohistochemistry of Dcx^+^Tshz1-GFP^+^ cells (arrows) in the PL at P7, P21, and P35. **g,h,** Quantification of the density of Dcx^+^Tshz1-GFP^+^ cells (**g**), and percentage of Dcx^+^ cells expressing Tshz1-GFP (**h**) at each age. *** = p < 0.001, **** = p < 0.0001. 1-way ANOVA with Holm-Sidak’s post-hoc test. Scale bars = 100 µm (**e, f** top row), 20 µm (**a, b, f** bottom row). PL: paralaminar nucleus; BLA: basolateral amygdala; ITCs: intercalated cell clusters; vEN: ventral endopiriform cortex.

**Extended Data Fig. 4:**
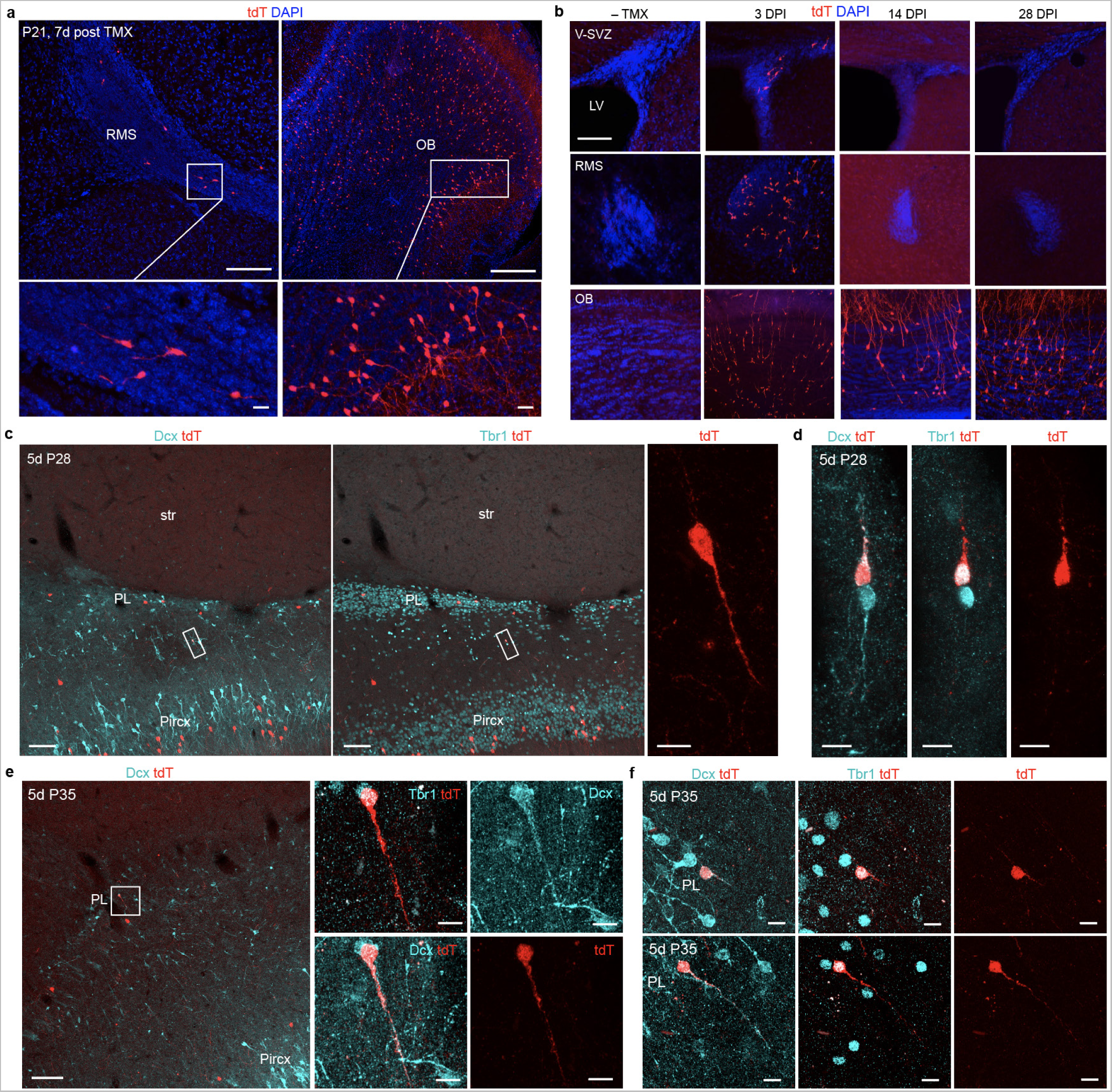
Dcx-CreER reporter mice expression pattern in the V-SVZ and PL. **a,** The RMS and OB contain tdT^+^ cells in an animal given TMX at P14 and examined at P21. **b,** TMX or no TMX control was given to adult animals and the V-SVZ, RMS, and OB were evaluated for tdT expression at 3, 14, or 28 days post injection (DPI). No TMX control mice did not have tdT^+^ cells in any of these regions. 3 DPI had tdT^+^ cells in all regions, whereas at 14 and 28 DPI the tdT cells were restricted to the OB. **c,d,** Mice given TMX on P21, 22, and 23 and sacrificed 5 DPI at P28 had tdT^+^ cells in the PL and piriform cortex. Overview image (**c**) shows the location of the cell shown in Fig. 4d. A pair of Dcx^+^Tbr1^+^ cells with one tdT^+^ with elongated processes extending opposite directions (**d**). **e,f,** Mice given TMX on P28, 29, and 30 and sacrificed 5 DPI at P35 had tdT^+^ cells in the PL and piriform cortex. A small tdT^+^ cell in the PL with one elongated process co-expressed Dcx (**e**). Similar cells could be observed in the PL and vEN co-expressing Dcx, Tbr1, and tdT (**f**). Scale bars = 200 µm (**a** top row) 100 µm (**b, c** left and middle, **e** left), 20 µm (**a** bottom row), 10 µm (**c** right, **d, e** right, **f, g**). Pircx: piriform cortex; RMS: rostral migratory stream; OB: olfactory bulb; V-SVZ: ventricular-subventricular zone; tdT: tdTomato; TMX: tamoxifen; DPI: days post injection.

**Extended Data Fig. 5:**
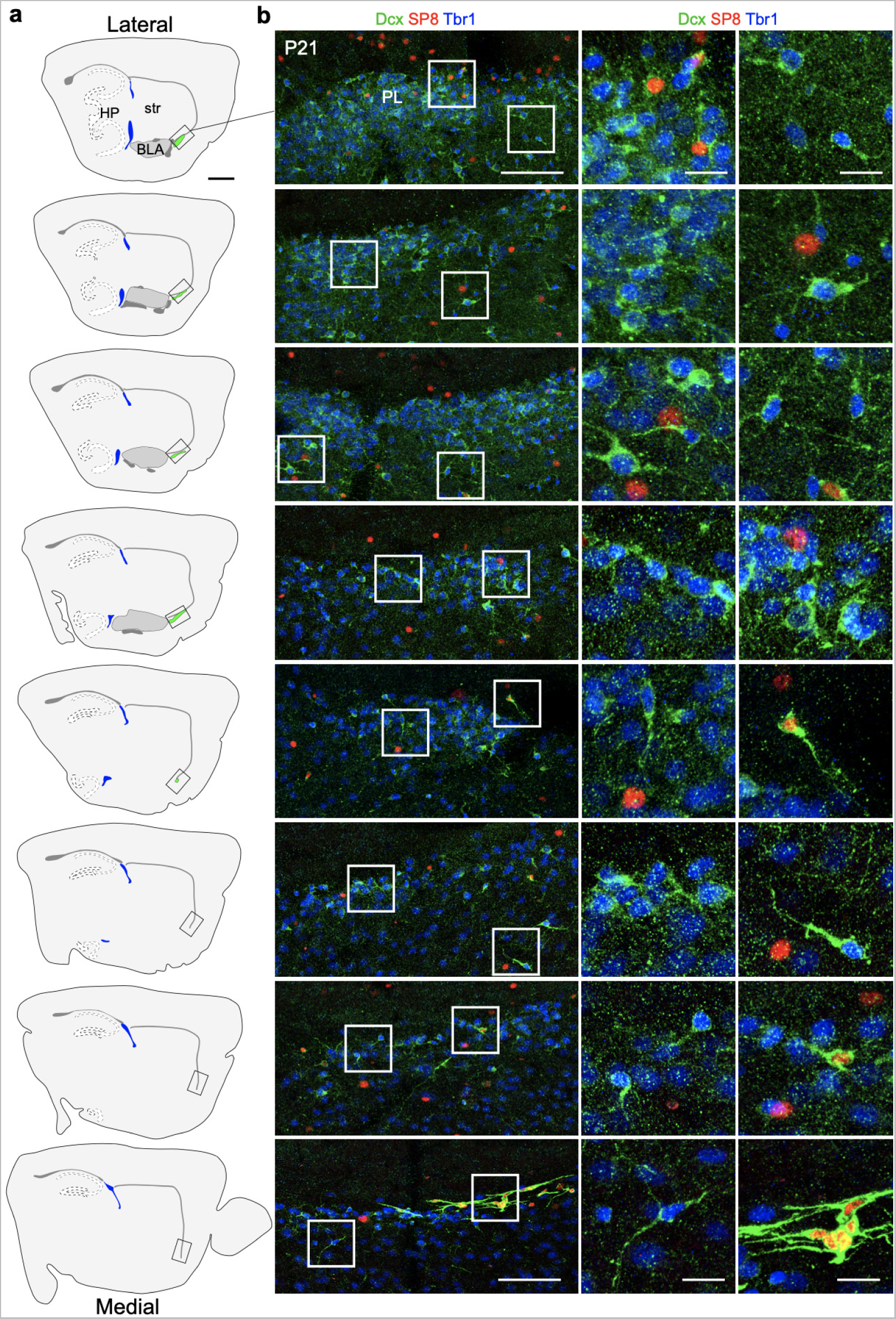
Migratory Tbr1^+^ cells in the PL are distinct from migratory Sp8^+^ cells in the RMS. **a,** Map of a lateral to medial series of sections each spaced by 200µm spanning the BLA to the edge of the RMS. The level of each immunostained section is indicated. **b,** Dcx^+^Tbr1^+^ cells or Dcx^+^Sp8^+^ cells at the level indicated in (**a**). The edge of the RMS is visible as a dense collection of Dcx^+^Sp8^+^ cells in the most medial section. Scale bars = 1 mm (a), 100 µm (b left), 20 µm (b middle and right). HP: hippocampus; PL: paralaminar nucleus, str: striatum.

**Extended Data Fig. 6:**
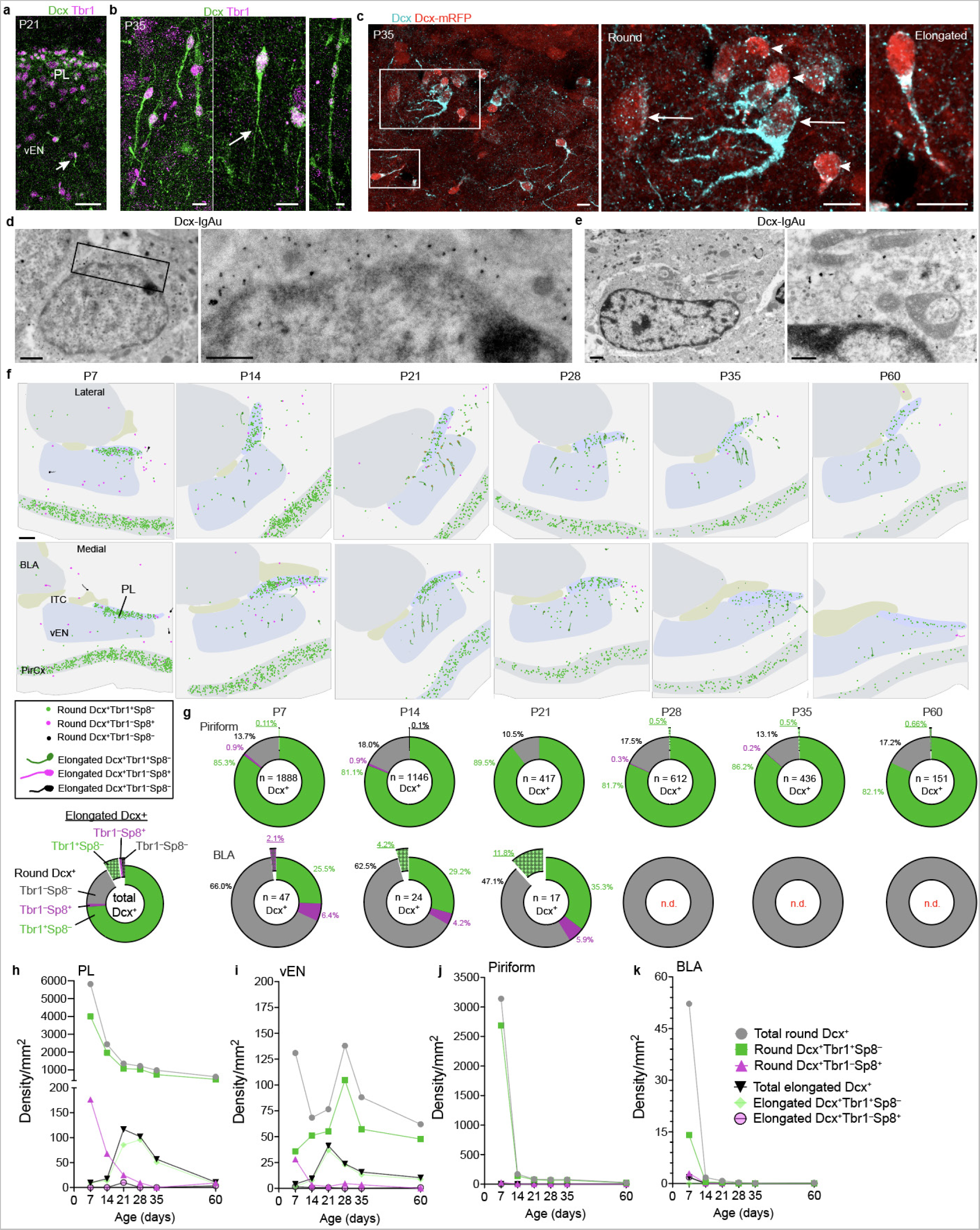
A wave of migration occurs between the PL and vEN during postnatal ages. **a,** At P21, Dcx^+^Tbr1^+^ cells can be seen with elongated processes in the vEN (arrow). **b,** At P35, similar Dcx^+^Tbr1^+^ cells are present as well as some with bifurcated processes (arrow) and in chains (right). **c,** Dcx-mRFP mice at P35 have Dcx^+^mRFP^+^ cells in the PL with small round (arrowheads) and larger more complex (arrows) morphologies. In the same field a Dcx^+^mRFP^+^ cell with an elongated morphology is also visible. **d,e,** Dcx-immunogold labeled cells in the PL at P21. Some labeled cells are small with few organelles, compact chromatin, dense immunogold labeling, and limited cytosol (**d**) while other cells have features of more mature cells such as more abundant organelles, more sparse immunogold labeling, and cell expansions (**e**). **f,** 1.5 mm square Maps of the location of round and elongated Dcx^+^ cells co-expressing either Tbr1 (green) or Sp8 (magenta) or neither (black) between P7 and P60. Additional are lateral (top row) or medial (bottom row) relative to each other. **g,** Quantification of the percentage of round and elongated Dcx^+^ cells co-expressing either Tbr1 (green) or Sp8 (magenta) or neither (black) between P7 and P60 in the Piriform cortex (top row) and BLA (bottom row). Elongated percentages are underlined and popped out of the chart. n.d. indicates that fewer than 5 Dcx^+^ cells were observed in the structure. Total number of Dcx^+^ cells evaluated in each region and age is indicated in the center of each chart. **h–k,** Density of round and elongated Dcx^+^ cells co-expressing either Tbr1 (green) or Sp8 (magenta) or all Dcx^+^ cells of each type (grey) between P7 and P60 in the PL (**h**), vEN (**i**), Piriform cortex (**j**), and BLA (**k**). Scale bars = 150 µm (**f**), 50 µm (**a**), 10 µm (**b, c**), 1 µm (**d** left, **e** left), 500 nm (**d** right, **e** right). PL: paralaminar nucleus; vEN: ventral endopiriform cortex; BLA: basolateral amygdala: ITC: intercalated cell cluster; PirCx: piriform cortex.

**Extended Data Fig. 7:**
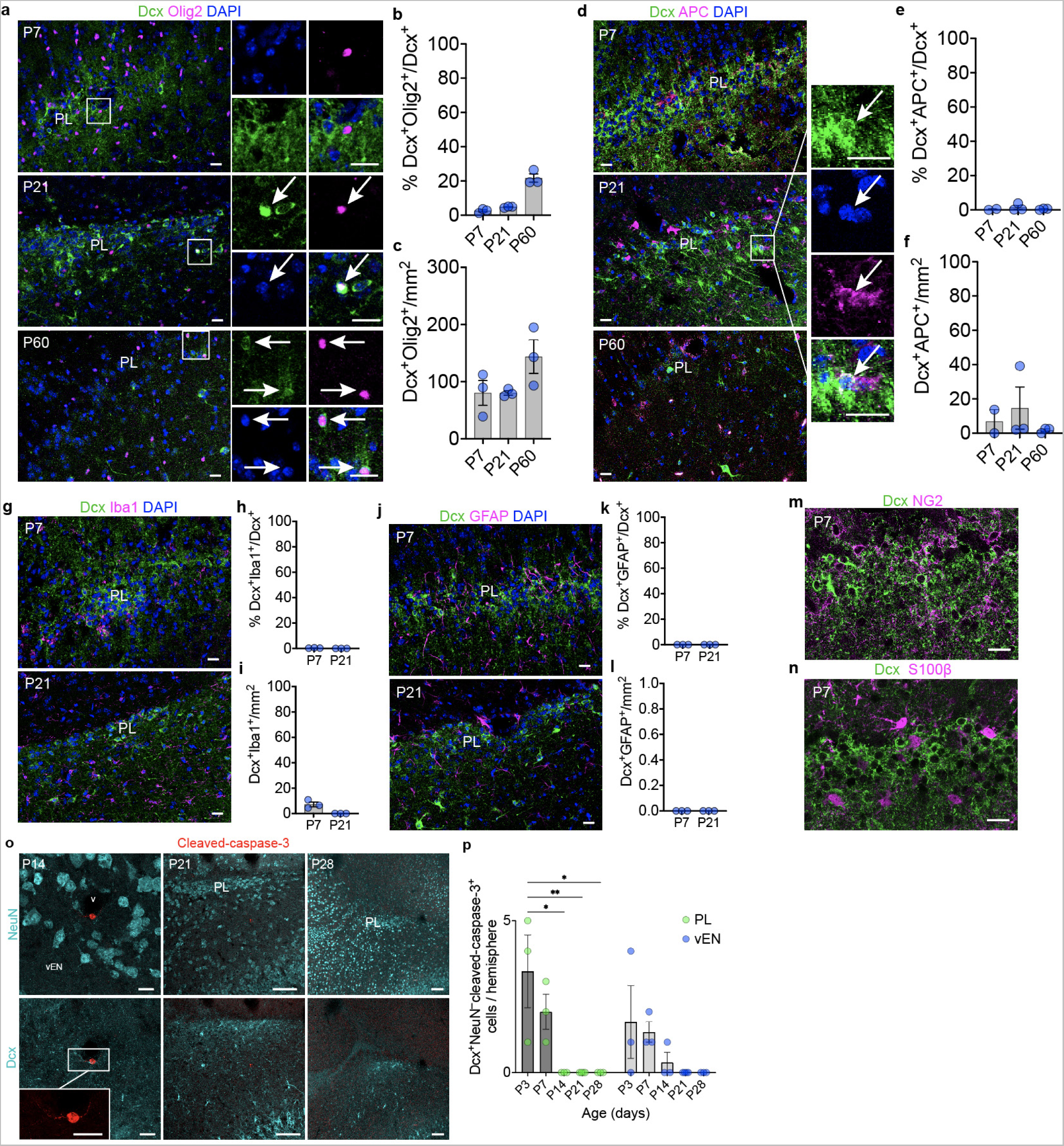
Dcx is expressed in the oligodendrocyte lineage postnatally. **a,** Immunostaining for Dcx^+^Olig2^+^ cells (arrows) in the PL at P7, P21, and P60. **b,c,** Quantification of the percentage of Dcx^+^ cells expressing Olig2 (**b**) and the density of Dcx^+^Olig2^+^ cells (**c**) at each age. **d,** Immunostaining for Dcx^+^APC^+^ cells (arrow) in the PL at P7, P21, and P60. **e,f,** Quantification of the percentage of Dcx^+^ cells expressing APC (**e**) and the density of Dcx^+^APC^+^ cells (**f**), at each age. **g,** Immunostaining for Dcx and Iba1 in the PL at P7 and P21. **h,i,** Quantification of the percentage of Dcx^+^ cells expressing Iba1 (**h**) and the density of Dcx^+^Iba1^+^ cells (**i**), at each age. **j,** Immunostaining for Dcx and GFAP in the PL at P7 and P21. **k,l,** Quantification of the percentage of Dcx^+^ cells expressing GFAP (**k**) and the density of Dcx^+^GFAP^+^ cells (**l**) at each age. **m,n,** Immunostaining for Dcx and NG2 (**m**) or S100β (**n**) in the PL at P7. **o,** Examples of staining for cleaved-caspase-3 in the PL and vEN at P14, P21, and P28; the same field of view is shown for different co-stains (top/bottom). One small peri-vascular Dcx^−^NeuN^−^ cell was observed (inset). **p,** Total number of Dcx^+^NeuN^−^cleaved-caspase-3^+^ cells per hemisphere in the PL and vEN between P3 and P28. * = p < 0.05, ** = p < 0.01. 2-way ANOVA with Holm-Sidak’s post-hoc test. Scale bars = 100 µm (**o** middle and right), 20 µm (**a, d, g, j, m, o** left). PL: paralaminar nucleus; vEN: ventral endopiriform cortex; v: blood vessel.

**Extended Data Fig. 8:**
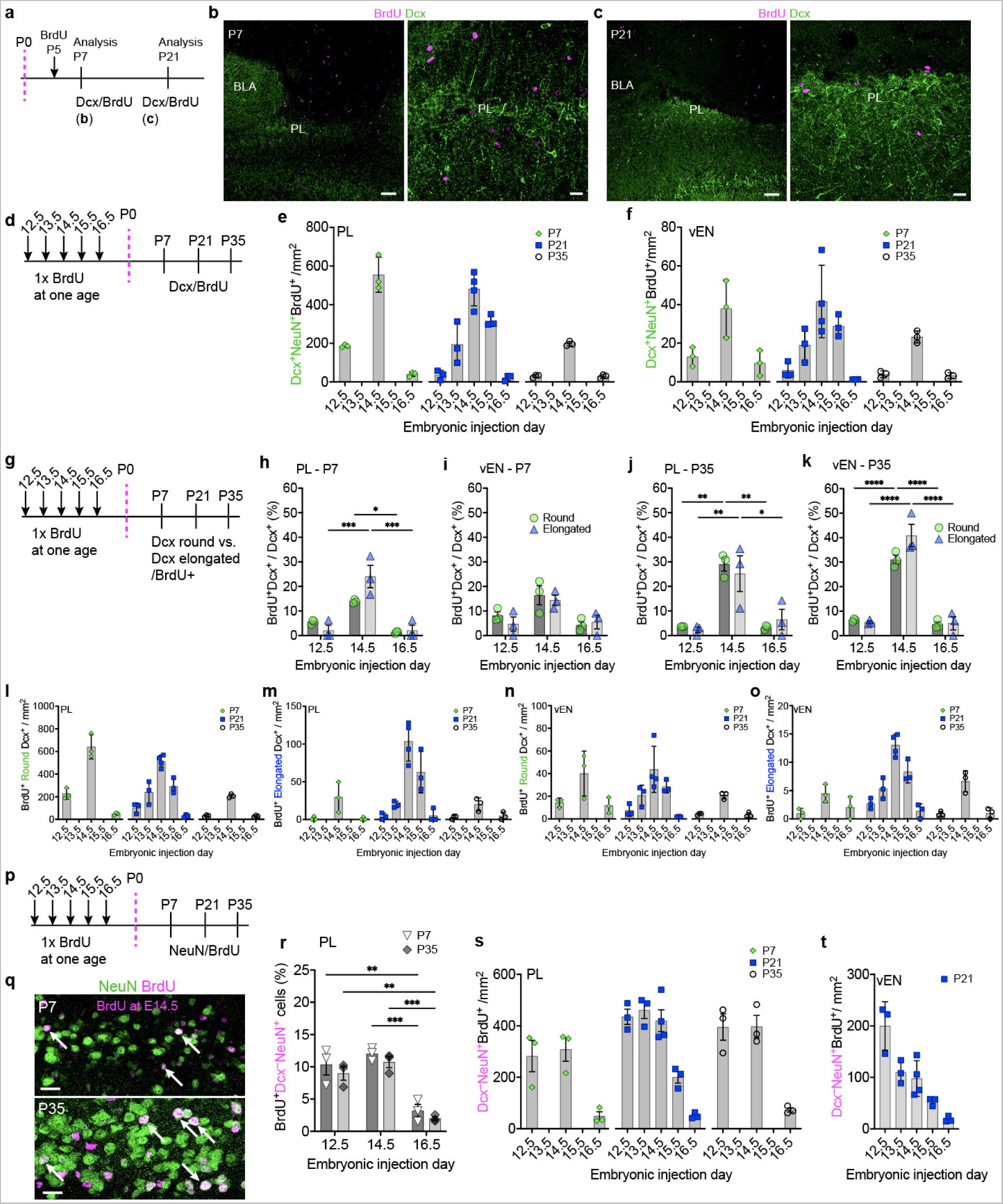
PL cells are born embryonically. a–c, BrdU was administered once at P5 and the PL was examined at either P7 (**b**) or P21 (**c**). No overlap between Dcx or BrdU was observed in the PL at either age. **d–f,** BrdU was given once to timed-pregnant mice between E12.5 – E16.5 and the density of Dcx^+^NeuN^+^ cells labeled with BrdU in the PL (**e**) or vEN (**f**) was examined in the offspring. **g–o**, BrdU was given once to timed-pregnant mice between E12.5 – E16.5 and the percentage (**h–k**) and density (**l–o**) of Dcx^+^ cells displaying either round or elongated morphology that co-labeled with BrdU in the PL (**h,j,l,m**) or vEN (**j,k,n,o**) was examined in the offspring. **p–t,** BrdU was given once to timed-pregnant mice between E12.5 – E16.5 and the percentage (**r**) and density (**s,t**) of Dcx^−^NeuN^+^ cells labeled with BrdU in the PL (**r,s**) or vEN (**t**) was examined in the offspring. * = p < 0.05, ** = p < 0.01, *** = p < 0.001, **** = p < 0.0001. 2-way ANOVA with Holm-Sidak’s post-hoc test. Scale bars = 100 µm (**b** left, **c** left), 20 µm (**b** right, **c** right, **h**). BLA: basolateral amygdala; PL: paralaminar nucleus; vEN: ventral endopiriform cortex.

**Extended Data Fig. 9:**
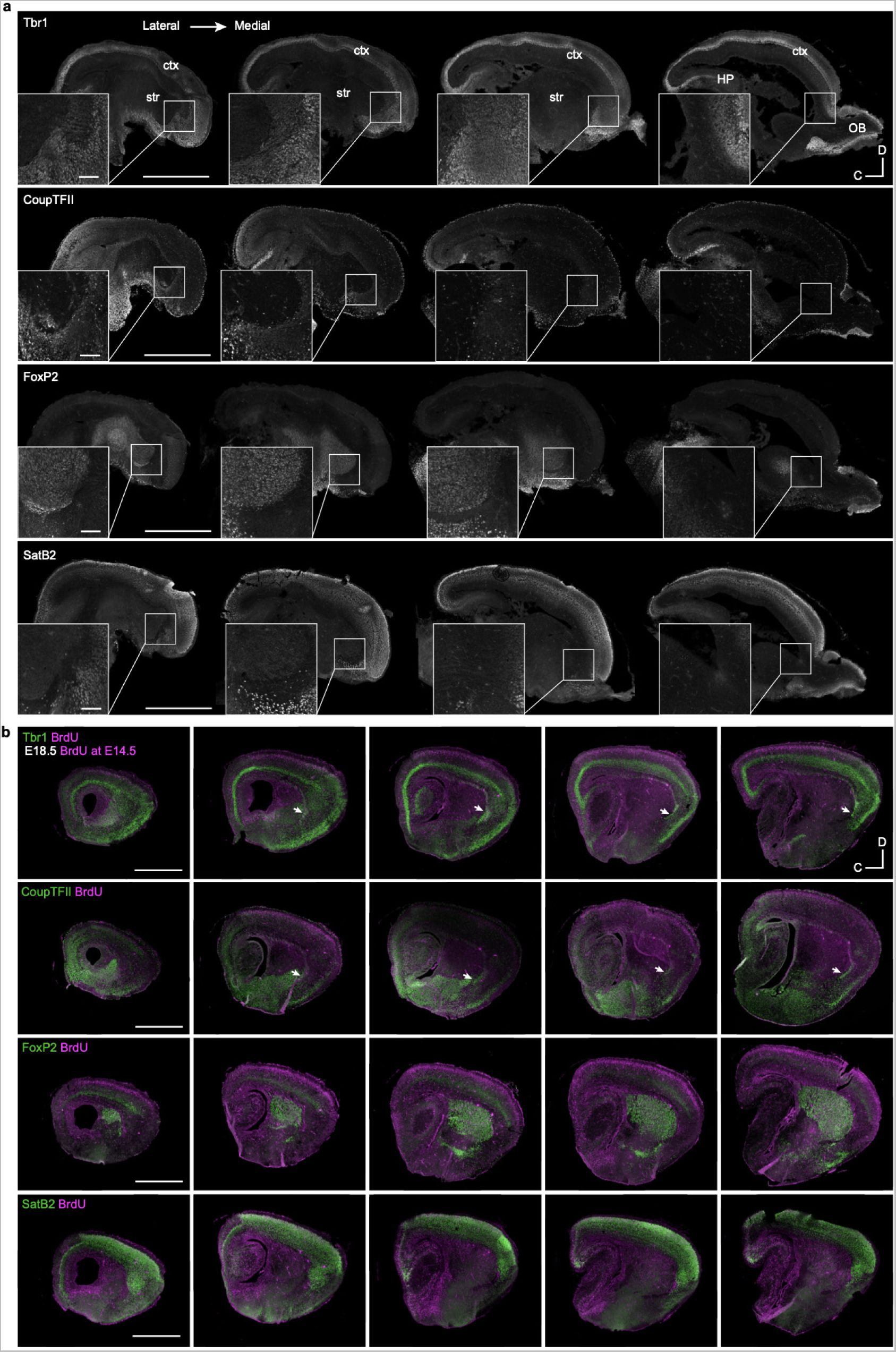
E14.5 BrdU labels a nucleus of Tbr1^+^CoupTFII^+^ cells during embryonic ages. **a,** E16.5 series of stains for Tbr1, CoupTFII, FoxP2, and SatB2. A dense nucleus of Tbr1^+^CoupTFII^+^ cells is not visible at this age. **b,** E18.5 series of stains for Tbr1, CoupTFII, FoxP2, and SatB2 in mice given one injection of BrdU at E14.5. A collection of Tbr1^+^CoupTFII^+^ cells (arrow) is visible anterior-ventral to the BLA and between cells expressing FoxP2 and SatB2. Scale bars = 1 mm (**a** overviews, **b**), 100 µm (**a** insets). ctx: cortex; HP: hippocampus; OB: olfactory bulb.

**Extended Data Fig. 10:**
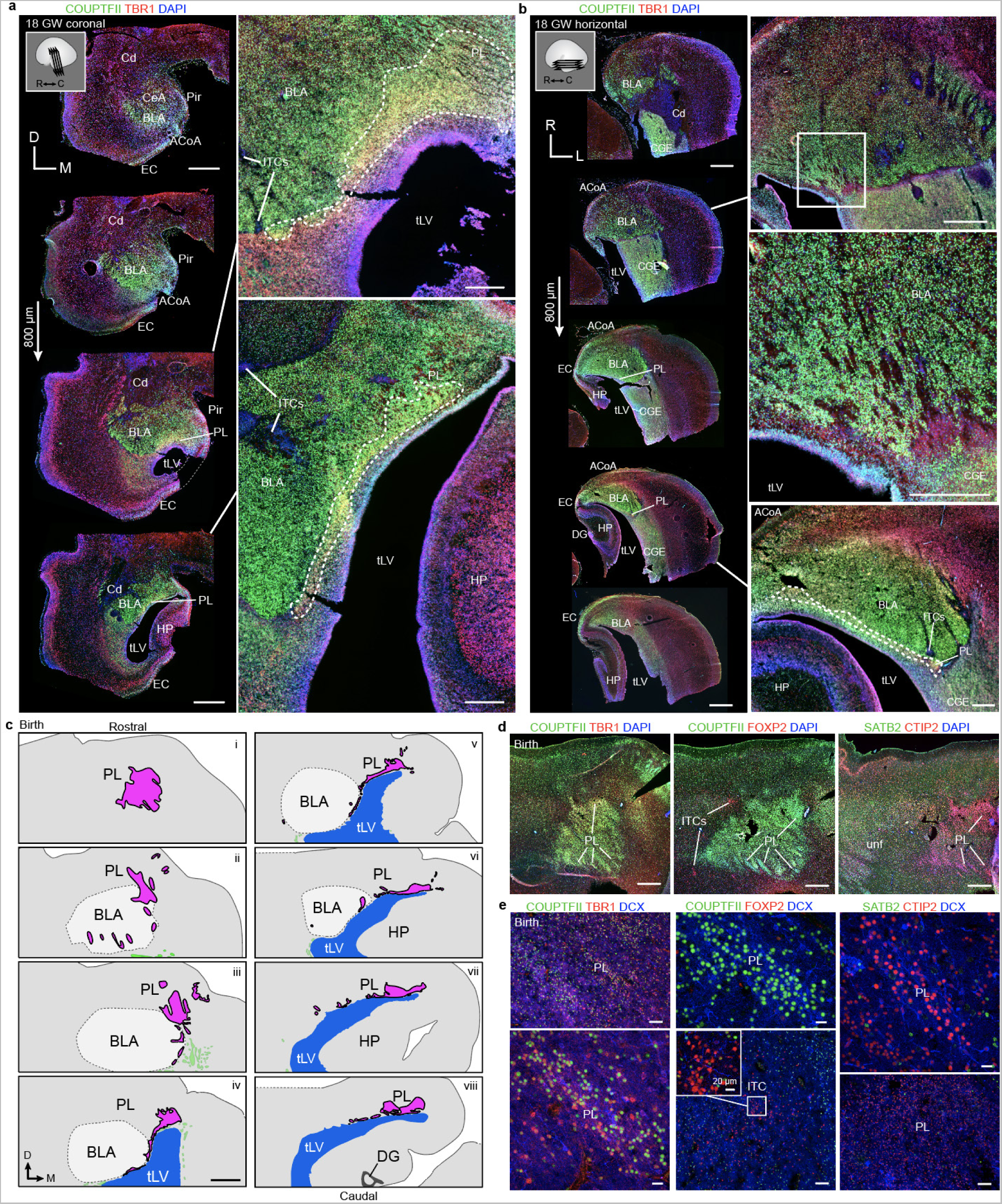
The human PL is COUPTFII^+^TBR1^+^CTIP2^+^SATB2^−^FOXP2^−^ and forms by 18 GW, extending rostrally. a, Human temporal lobe at 18 GW coronal sections from rostral to caudal stained for COUPTFII and TBR1. Two sectioning levels adjacent to the temporal lobe lateral ventricle (tLV) include an anatomical region with COUPTFII and TBR1 co-localized cells with the shape and location of the PL (insets dotted lines). **b,** Human temporal lobe at 18 GW horizontal sections from dorsal to ventral stained for COUPTFII and TBR1. Top magnified section displays some striated features typical of the PL, but limited COUPTFII and TBR1 co-localization. Bottom magnified section contains a thin COUPTFII^+^TBR1^+^ region (inset dotted lines). **c,** Maps of the location of the PL in the human amygdala at birth in coronal sections from rostral to caudal. Section (i) is at the level of immunostaining in (**d**), section (ii) is at the level in (**Fig. 8j–l**). Sections iii–viii are reproduced with additional details from Sorrells *et al.* 2019. **d,** Immunostaining for COUPTFII and TBR1, COUPTFII and FOXP2, or SATB2 and CTIP2 at birth. **e,** High magnification of immunostaining in (d) showing COUPTFII^+^TBR1^+^CTIP2^+^ cells in the PL that are FOXP2^−^ and SATB2^−^. The ITCs contain FOXP2^+^COUPTFII^−^ cells (inset). Scale bars = 2 mm (**a** left, **b** left, **c**), 1 mm (**b** top right, **d**), 500 µm (**a** right, **b** right middle and bottom), 100 µm (**e** top left and bottom middle), 20 µm (**e** bottom left top middle and right). ACoA: amygdala cortical area; BLA: basolateral amygdala; CeA: central amygdala; Cd: caudate; CGE: caudal ganglionic eminence; DG: dentate gyrus; EC: entorhinal cortex; GW: gestation week; HP: hippocampus; ITC: intercalated cell cluster; LPL: lateral paralaminar nucleus: MPL: medial paralaminar nucleus: PL: paralaminar nucleus; Pir: piriform cortex; tLV: temporal lobe lateral ventricle; unf: uncinate faciculus.

## Materials and Methods

### Mice

Male and female C57Bl/6J (Jackson labs, Strain #:000664) wildtype mice used in this study were socially housed (up to 4 mice/cage) on a 12 hour day/night cycle with ad libitum food and water in a pathogen-free facility. All experiments were conducted in accordance with approved protocols by the University of Pittsburgh Institutional Animal Care and Use Committee (IACUC) and following NIH guidelines for animal research. All mice used in this study were healthy and were only used in the manner described for each experiment and did not undergo previous experimental procedures. A transgenic construct using doublecortin (Dcx) BAC clone (GENSAT1-BX214) to express CreER^TM^ recombinase^91^ was injected into fertilized C57BL/6J eggs. Founder line was established by expression testing via subsequent crosses to a Rosa-LSL-tdTomato reporter^92^. Allele specific genotyping of transgenic mice was performed by PCR with primers (Cre: 5′-CTG CCA CCA GCC AGC TAT CAA CTC-3′) and (ER: 5′-CTC AGC ATC CAA CAA GGC ACT GAC-3′). Tshz1-GFP mice were a gift from Kenneth Campbell (Cincinnati Children’s Hospital) and Dcx-mRFP mice were a gift from Qiang Lu (City of Hope).

### 5-bromo-2-deoxyuridine (BrdU) administration

To generate timed pregnant mice, C57Bl/6J females were paired with a male for 24 hrs and then separated. Pregnant mice were given i.p. injections of 50 mg/kg BrdU (Thermo Scientific, catalog # H27260-06) dissolved in sterile saline at specific embryonic time points, considering the day post pairing as E0.5. For postnatal BrdU experiments, C57Bl/6J mice were given i.p. injections of 50 mg/kg BrdU singly or once-daily for seven days.

### Tamoxifen (TMX) administration

Cre positive Dcx-CreER^TM^::Rosa-LSL-tdTomato mice were given intraperitoneal injections of 10 mg/kg TMX (Sigma Aldrich, catalog # T5648) dissolved in sunflower oil (Sigma Aldrich, catalog # S5007) once daily for three days starting at either P21 or P28. Mice were then collected either 3, 5, or 12 days after the final injection.

### Mouse tissue immunostaining

Mice were anesthetized using an overdose of isoflurane and were transcardially perfused with 1x phosphate buffered saline (PBS) followed by 4% paraformaldehyde (PFA). Brains were postfixed in 4% PFA overnight and sunk in 30% sucrose in 1X PBS solution for 24 hr prior to sectioning on a freezing-sliding microtome at 50 µm. Sections were stored in 1X PBS + 0.05% sodium azide until staining. Unless otherwise indicated, stains were performed on a series of 1 in every 4 sections (200 µm apart). For most staining, a standard protocol was performed: sections were washed in 1x PBS + 0.2% triton (PBST) for 10 min, treated with (0.5%) sodium borohydride for 15 min, washed 5 times for 10 min per wash with 1x PBS followed by 1.5 hr incubation in 10% normal donkey serum (NDS) in PBST (block solution). Sections were washed 3x for 10 min in PBST followed by incubation in primary antibody (**Supplementary Table 4**) in block solution overnight (16-24 hrs) at 4 °C. Sections were then washed with PBST 5 times for 10 min per wash before being incubated in secondary antibodies (**Supplementary Table 4**) and DAPI (1:5000 dilution, Fisher, catalog # ICN15757401) in block solution for 90 min at room temperature. The sections were then washed a final time with PBST 5 times for 10 min per wash and then mounted with Fluoromount-G (SouthernBiotech, catlog # OB100-01).

For postnatal BrdU co-staining, sections were washed and then treated with 2N HCl for 30 min at 37 °C, then washed 5 times for 10 min per wash with 1X PBST. Sections were then stained with BrdU primary antibody and other co-stains according to the standard staining protocol above. For embryonic sections, 20 µm cryostat sections (Cryostar NX50, ThermoFisher) were mounted to Superfrost Plus slides (Fisher, catalog # 12-544-2) and stored at −80 °C until use. BrdU co-staining in embryonic tissue was performed by first staining for all other primary antibodies according to the standard staining protocol. The tissue was then treated with 4% PFA for 10 min, followed by incubation in 2N HCl for 20 min at 37 °C, washed 5 times with 10 min per wash, and then stained with BrdU primary antibodies overnight followed by detection using the standard protocol above.

### Anatomical boundaries of the developing and adult mouse paralaminar nucleus

The PL is most readily visualized in the sagittal sectioning plane due to its elongation across the rostral-caudal axis. In sagittal sections, at its most lateral edge, the PL is a horn-shaped nucleus of small, densely packed cells, anterior and ventral to the basal lateral amygdala. In more medial planes, the PL is located more anteriorly and becomes separated from the BLA. In the 2011 Allen sagittal adult mouse brain atlas (Available from mouse.brain-map.org. Allen Institute for Brain Science (2011)), the PL is visible in the P56 Nissl staining sagittal series beginning on section three and ending on section seven. This region is varyingly indicated as belonging to one or multiple neighboring structures depending on the section level; however, as we describe in the text it is molecularly and cellularly distinctive as its own region.

### Unbiased Stereology

Stereological quantifications in the PL and vEN were performed with the optical fractionator probe using Steroinvestigator (MBF Biosciences) on a Leica DM4 microscope. Using a 10x/0.45 NA objective, the boundaries of the PL and vEN were delineated based on the cytoarchitecture visible from DAPI and NeuN staining, which remain constant across the ages examined (e.g. **Fig. 2j, Extended Data Fig. 3e**). DAPI^+^, Dcx^+^ and NeuN^+^ cells were counted using a 63x/1.40 NA oil-immersion objective. Section thickness was measured at every counting site. Five sex-balanced mice were counted at P7, P14, P21, P35, and P60. Five 50 µm sections, each spaced by 200 µm, were counted, which covered the extent of PL and vEN. To minimize section edge damage, a new disposable cutting blade was used for each brain during sectioning on a frozen sliding microtome. Mice in which a section was missing or an ROI was damaged were excluded from counts. To ensure a minimum of 150 objects were counted for each probe run across different sizes of objects and ages, the frame and grid sizes for each probe type were optimized (see **Supplementary Table 1** for a complete list of sampling parameters). For ages P21–P60, all probes were counted with a dissector height of 15µm with 5µm guard zones. Tissue from mice at younger ages, P7 and P14, exhibited greater z-collapse, so we used a dissector height of 13 µm and 3 µm guard zones. The unique point on each counted object (its leading edge) that was used for inclusion/exclusion criteria was the boundary of the soma. Objects were counted if they intersected with a inclusion line of the counting frame or were within the counting frame entirely. The end of the soma and beginning of a process (which was not used for counting) was defined by the inflection point at which the process began to taper away from the soma. All probe runs included in the study had a Gunderson coefficient of error (CE) (Gundersen, 1999) below 0.1. The average Gunderson CEs for each probe and region were: PL DAPI = 0.0757, PL NeuN CE = 0.0590, PL Dcx CE = 0.0667, vEN DAPI CE = 0.0670, vEN NeuN CE = 0.0623, and vEN Dcx CE = 0.0783.

### Marker co-expression quantification

For all marker co-expression quantifications, five 50 µm sections of tissue evenly spaced by 200 µm (1 out of every 4 sections) were evaluated. For Dcx, NeuN and transcription factor co-localization quantifications we used a Leica TCS SP8 confocal microscope (20x/0.75 NA objective) to collect system optimized z-stacks of the entire tissue z-height of the PL. Co-expression of the specified transcription marker and Dcx^+^NeuN^+^ and Dcx-NeuN^+^ cells in the PL was quantified using ImageJ (Rasband, W.S., ImageJ, U. S. National Institutes of Health, Bethesda, Maryland, USA, https://imagej.nih.gov/ij/, 1997-2018). Individual channels were merged and accessed across each frame of the z stack. For Dcx-CreER^TM^::Rosa-LSL-tdTomato quantifications, we used a Leica TCS SP8 confocal microscope (10x/0.40 NA objective) to collect system optimized z-stacks of the entire tissue z-height of the PL and quantified co-expression using ImageJ. For BrdU quantifications, we used StereoInvestigator (MBF Biosciences) connected to a Leica DM4 microscope. Boundaries of the PL and vEN were traced using a 10x/0.45 NA objective and co-localization was assessed across the z-height of the section with a 63x/1.40 NA oil-immersion objective. Dcx+ cells were classified as either round or elongated, as described below.

### Cell morphology reconstructions and mapping

To map the location of Dcx^+^ neurons with round or elongated morphology, cells were counted and traced using Neurolucida (MBF Biosciences) on a Leica DM4 microscope with a 10x/0.45 NA objective (for ROI delineation) and a 63x/1.40 NA oil-immersion objective (for cell counting and cell tracing). Within a 1.5 mm square region centered on the PL, we traced the outline of the BLA, ITCs, vEN, layer 2 of the piriform cortex, and PL. The location of every Dcx^+^ cell within that square was mapped together with information about co-expression and morphology of each cell. A Dcx^+^ cell was classified as elongated and traced if its soma was fusiform and also had one simple (unbranched) linear projection. A Dcx^+^ cell was classified as round if its soma was spherical and/or if it had more than one process and/or if it had a single process with multiple branches.

### Transmission electron microscopy and immunogold labeling

Mice (P14, P21 and P28) were deeply anesthetized and perfused with 0.9% saline followed by 2% paraformaldehyde (PFA) and 2.5% glutaraldehyde (EMS, Hatfield, PA, USA) in 0.1 M phosphate buffer (PB). Brains were dissected and post-fixed overnight at 4 °C in the same fixative solution and subsequently, 200 µm sagittal sections were prepared using a Leica VT1200S vibratome (Leica Microsystems GmbH, Heidelberg, Germany). Slices were further post-fixed in 2% osmium tetroxide in 0.1 M PB for 1.5 hours at room temperature, washed in deionized water, and partially dehydrated in 70% ethanol. Samples were then incubated in 2% uranyl acetate in 70% ethanol in the dark for 2.5 h at 4 °C. Brain slices were further dehydrated in ethanol followed by propylene oxide and infiltrated overnight in Durcupan ACM epoxy resin (Fluka, Sigma-Aldrich, St. Louis, USA). The following day, the samples were transferred to fresh resin and were cured for 72 h at 70 °C. Following resin hardening, semithin sections (1.5 µm) were obtained with a diamond knife using a Ultracut UC7 ultramicrotome (Leica). Sections were mounted onto glass microscope slides and lightly stained with 1% toluidine blue. After identification of the PL by light microscopy, selected semithin sections were glued with Super Glue-3, Loctite (Henkel, Düsseldorf, Germany) to resin blocks and subsequently detached from the glass-slides by repeated freezing (in liquid nitrogen) and thawing. Ultra-thin sections (60–80 nm) were obtained with the ultramicrotome from detached semi-thin sections, and further stained with lead citrate (Reynolds’ solution). The samples were examined with a FEI Tecnai G2 Spirit transmission electron microscope at 80 kV (FEI Europe, Eindhoven, Netherlands) equipped with a Morada CCD digital camera (Olympus Soft Image Solutions GmbH, Münster, Germany).

For pre-embedding immunogold stains, samples were fixed with 4% PFA in 0.1 M PB and cut into 50 µm sagittal sections using a vibratome. Pre-embedding immunogold stainings were performed (see **Supplementary Table 4**) as previously described^93^. Sections were contrasted with 1% osmium tetroxide, 7% glucose in 0.1 M PB and embedded in Durcupan epoxy resin. Subsequently, 100-150 serial semithin sections (1.5 µm) were obtained with the ultramicrotome, mounted onto glass-slides, and stained with 1% toluidine blue. Selected semi-thin sections were processed as described above to obtain ultrathin sections prior to examination by TEM.

### iDISCO staining and light sheet microscopy

For whole mount imaging, the iDisco tissue clearing method was used^94^. Samples were perfused in 4% paraformaldehyde overnight at 4 °C, and then washed in PBS. Prior to immunostaining, tissue was dehydrated using increasing concentrations of methanol in distilled water (20%, 40%, 60%, 80%, 100%, 100%, 1 hour each at RT), bleached overnight in 5% H2O2 in 100% methanol at 4 °C, and then gradually rehydrated (80%, 60%, 40%, 20% methanol, then PBS with 0.2% Triton, 1 hour each at RT). Bleached samples were incubated in permeabilization solution (PBS with 0.2% Triton, 20% DMSO, 0.3M glycine) and then in blocking solution (PBS with 0.2% Triton, 10% DMSO, 6% Normal Donkey Serum), each for 2 days at 37 °C. Primary antibody incubation was performed in PTwH (PBS with 0.2% Tween-20 and 10 µg/ml heparin) + 5% DMSO and 3% Normal Donkey Serum, 4 days at 37 °C. Samples were washed in PTwH over 24 hours at RT (5-6 washes of increasing duration), before incubation in secondary antibody in PTwH + 3% Normal Donkey Serum, 4 days at 37 °C. Another series of 5-6 washes in PTwH over 24 hours at RT were performed before proceeding to clearing steps.

Clearing was performed by dehydrating samples in increasing concentrations of methanol in distilled water (20%, 40%, 60%, 80%, 100%, 1 hour each at RT), then incubating in a solution of 66% dichloromethane (DCM, Sigma 270997) in methanol for 3 hours at RT. 2 washes of 100% DCM (15 minutes each at RT) were performed before placing samples in dibenzyl ether (DBE, Sigma 108014) for clearing and imaging. Samples were imaged in DBE via light sheet microscopy (Ultramicroscope II, Miltenyi Biotec) equipped with an sCMOS camera (Andor NEO 5.5) and a 2x Olympus objective lens. Cleared samples were stored long-term in DBE in glass containers in the dark. Light sheet datasets were imported into Imaris 9.1 (Bitplane) for 3D visualization. No masking was performed. Following automatic rendering using the ‘MIP’ setting, videos of 3-D datasets were generated using the ‘animation’ tool.

### Time-lapse microscopy and migratory neuron tracking

For time-lapse microscopy, P0.5-P3 Dcx mRFP mice were anesthetized in wet ice before being cleaned with 70% ethanol and decapitated with sterilized scissors. The brain was then extracted and placed in chilled ACSF (glucose 25 mM, NaCl 125, KCl 2.5 mM, MgCl2 1 mM, CaCl2 2 mM, NaHCO3 2.5 mM, Na2HPO4 1.25 mM) or HBSS (Gibco, catalog # 14175079) that had been bubbled with carbogen (95% O2 and 5% CO2) for 30 minutes while on wet ice. Brains were embedded in blocks with 3.5% low melting point agar (Fisher, catalog # 16-520-050) heated to 37 °C. After the agar had solidified, 350 μm sagittal sections were prepared using a vibratome (Leica VT 1200S) with carbogen-bubbled ACSF or HBSS in the sectioning chamber that was externally packed with wet ice. Sections were transferred onto sterile membrane culture inserts (Millicell-CM, Millipore; 0.4 μm pore size, 30 mm diameter). These were placed into either a glass-bottom coverslip petri dish or a six-well plate containing pre-incubated slice culture medium (1.5-2 mL per well) that consisted of 66.6% BME with phenol red, 25% HBSS, 5% FBS non-heat inactivated, 1.3% of 33% glucose, 1% P/S/L-glut, 1% Glutamax (L-glut 200 mM), and 1% N2. Cultures were imaged on the microscope in a humidified incubator (okoLab Transparent Cage Incubator DMi8, product # 158206046) with a heated and humidified gas mixture (5% O2, 5% CO2, at 37 °C). Imaging was every 20-30 min using a Leica Sp8 confocal resonance scanner and a 10x/0.40 NA objective with 2-3x digital zoom.

Traces of migrating neurons were generated using the ImageJ manual tracking feature. The coordinates of the soma of each migrating neuron was recorded for each time frame it was visible within the optical section. Those coordinates were then used to calculate each neuron’s total distance traveled and total displacement.

### Human tissue collection

Twenty-one post-mortem specimens were examined for this study (**Supplementary Table 3**). Tissue was collected with previous patient consent in strict observance of the legal and institutional ethical regulations in accordance with each participating institution: 1. In accordance with institutional guidelines and study design approval by the Committee for Oversight of Research and Clinical Training Involving Decedents (CORID) at the University of Pittsburgh. 2. Specimens collected at the University of Pittsburgh Medical Center (UPMC) had IRB approved research informed consents along with HIPAA authorizations signed by parents or responsible guardians. 3. The University of California, San Francisco (UCSF) Committee on Human Research. Protocols were approved by the Human Gamete, Embryo and Stem Cell Research Committee (Institutional Review Board) at UCSF. For infant cases, when the brain is at full term (37 to 40 gestational weeks), we refer to this as “birth”. We collected tissue blocks from the temporal lobe, anteriorly from the amygdaloid complex to the posterior end of the inferior horn of the lateral ventricle. Samples were either flash frozen or fixed in 4% paraformaldehyde (PFA) or 10% formalin for >24h. Brains were cut into ~1.5 cm blocks, cryoprotected in a series of 10%, 20%, and 30% sucrose solutions, and then frozen in an embedding medium, OCT. Blocks of the medial temporal lobe were cut into 20 micron sections on a cryostat (Leica CM3050S) and mounted on glass slides for immunohistochemistry.

### Human tissue immunofluorescence

Frozen slides were allowed to equilibrate to room temperature for 3 hours and rehydrated in PBS prior to antigen retrieval at 95°C in 10 mM Na citrate buffer, pH=6.0. Slides were washed with TNT buffer (0.05% TX100 in PBS) for 10 minutes, placed in 1% H2O2 in PBS for 45 minutes and then blocked with TNB solution (0.1 M Tris-HCl, pH 7.5, 0.15 M NaCl, 0.5% blocking reagent from Akoya Biosciences) for 1 hour. Slides were incubated in primary antibodies overnight at 4°C (**Supplementary Table 4**) and in biotinylated secondary antibodies (Jackson Immunoresearch Laboratories) for 2.5 hours at room temperature. All antibodies were diluted in TNB solution. For most antibodies, the conditions of use were validated by the manufacturer (antibody product sheets). When this information was not provided, we performed control experiments, including no primary antibody (negative) controls and comparison to mouse staining patterns. Sections were then incubated for 30 min in streptavidin-horseradish peroxidase that was diluted (1:200) with TNB. Tyramide signal amplification (PerkinElmer) was used for some antigens. Sections were incubated in tyramide-conjugated fluorophores for 5 minutes at the following dilutions: Fluorescein: 1:100; Cy3: 1:100; Cy5: 1:100. After several PBS rinses, sections were mounted in Fluoromount G (Southern biotech) and coverslipped. Staining was conducted in technical triplicates prior to analysis.

### Statistics

Statistical significance was defined as * = p < 0.05, ** = p < 0.01, *** = p < 0.001, **** = p < 0.0001. One way or two way analysis of variance with Holm-Sidak post hoc tests was performed using GraphPad Prism (v.9). Quantifications are shown as mean ± SEM.

## Supplementary Files

Supplementary Table 1: Stereology Parameters

Supplementary Table 2: Stereology Results

Supplementary Table 3: Human Case Table

Supplementary Table 4: Antibodies

Supplementary Movie 1: P7 lightsheet imaging

Supplementary Movie 2: P21 lightsheet imaging

Supplementary Movie 3: PL migration

