## Supplementary Table 1 for "Mouse paralaminar amygdala excitatory neurons migrate and mature during adolescence"

**Supplementary Table 1: Stereology Parameters**

| ROI | Age | Probe | Grid size (µm) | Frame size (µm) | Guard zone (top-bottom µm) | Dissector height (µm) |
|---|---|---|---|---|---|---|
| PL | P7 | DAPI | 50x50 | 15x15 | 3-3 | 13 |
| PL | P7 | Neun | 50x50 | 25x25 | 3-3 | 13 |
| PL | P7 | Dcx | 40x40 | 25x25 | 3-3 | 13 |
| PL | P14 | DAPI | 65x65 | 15x15 | 3-3 | 13 |
| PL | P14 | Neun | 57x57 | 25x25 | 3-3 | 13 |
| PL | P14 | Dcx | 58x58 | 25x25 | 3-3 | 13 |
| PL | P21 | DAPI | 65x65 | 15x15 | 5-5 | 16 |
| PL | P21 | Neun | 57x57 | 25x25 | 5-5 | 16 |
| PL | P21 | Dcx | 57x57 | 25x25 | 5-5 | 16 |
| PL | P28 | DAPI | 65x65 | 15x15 | 5-5 | 16 |
| PL | P28 | Neun | 57x57 | 25x25 | 5-5 | 16 |
| PL | P28 | Dcx | 57x57 | 35x35 | 5-5 | 16 |
| PL | P35 | DAPI | 65x65 | 15x15 | 5-5 | 16 |
| PL | P35 | Neun | 57x57 | 25x25 | 5-5 | 16 |
| PL | P35 | Dcx | 57x57 | 35x35 | 5-5 | 16 |
| PL | P60 | DAPI | 65x65 | 15x15 | 5-5 | 16 |
| PL | P60 | Neun | 57x57 | 25x25 | 5-5 | 16 |
| PL | P60 | Dcx | 57x57 | 40x40 | 5-5 | 16 |
| vEN | P7 | DAPI | 110x110 | 20x20 | 3-3 | 13 |
| vEN | P7 | Neun | 110x110 | 40x40 | 3-3 | 13 |
| vEN | P7 | Dcx | 110x110 | 80x80 | 3-3 | 13 |
| vEN | P14 | DAPI | 115x115 | 20x20 | 3-3 | 13 |
| vEN | P14 | Neun | 120x120 | 30x30 | 3-3 | 13 |
| vEN | P14 | Dcx | 110x110 | 80x80 | 3-3 | 13 |
| vEN | P21 | DAPI | 120x120 | 20x20 | 5-5 | 16 |
| vEN | P21 | Neun | 120x120 | 45x45 | 5-5 | 16 |
| vEN | P21 | Dcx | 110x110 | 45x45 | 5-5 | 16 |
| vEN | P28 | DAPI | 110x110 | 20x20 | 5-5 | 16 |
| vEN | P28 | Neun | 120x120 | 30x30 | 5-5 | 16 |
| vEN | P28 | Dcx | 110x110 | 60x60 | 5-5 | 16 |
| vEN | P35 | DAPI | 120x120 | 20x20 | 5-5 | 16 |
| vEN | P35 | Neun | 120x120 | 35x35 | 5-5 | 16 |
| vEN | P35 | Dcx | 110x110 | 60x60 | 5-5 | 16 |
| vEN | P60 | DAPI | 140x140 | 20x20 | 5-5 | 16 |
| vEN | P60 | Neun | 140x140 | 35x35 | 5-5 | 16 |
| vEN | P60 | Dcx | 120x120 | 80x80 | 5-5 | 16 |
