## Supplementary Table 2 for "Mouse paralaminar amygdala excitatory neurons migrate and mature during adolescence"

**Supplementary Table 2: Stereology Results**

| Age | Sex | Region | Total Area (µm²) | Total Estimated Population (DAPI) | Total NeuN Estimated Population | Total Dcx Estimated Population | Measured volume (µm³) | CE Gundersen DAPI | CE Gundersen NeuN | CE Gundersen Dcx | Mean measured tissue thickness (µm) |
|---|---|---|---|---|---|---|---|---|---|---|---|
| P7 | F | PL | 378985 | 22021 | 9262 | 11219 | 75796900 | 0.06 | 0.04 | 0.04 | 18.257 |
| P7 | M | PL | 374135 | 21931 | 8448 | 11636 | 74827000 | 0.06 | 0.05 | 0.04 | 18.299 |
| P7 | M | PL | 292735 | 21982 | 8529 | 12813 | 58547000 | 0.07 | 0.06 | 0.05 | 19.1 |
| P7 | M | PL | 303443 | 21802 | 8437 | 11228 | 60688600 | 0.06 | 0.05 | 0.04 | 18.997 |
| P7 | F | PL | 356289 | 22334 | 8661 | 10971 | 71257800 | 0.05 | 0.05 | 0.04 | 17.86 |
| P14 | M | PL | 368053 | 21590 | 8898 | 9292 | 73610500 | 0.07 | 0.06 | 0.06 | 18.3 |
| P14 | F | PL | 367182 | 20403 | 8148 | 9522 | 73436400 | 0.07 | 0.06 | 0.06 | 19.003 |
| P14 | M | PL | 323920 | 19096 | 8543 | 10791 | 64783900 | 0.08 | 0.07 | 0.06 | 20.398 |
| P14 | F | PL | 310365 | 21507 | 8490 | 10384 | 62073000 | 0.08 | 0.06 | 0.06 | 20.635 |
| P14 | F | PL | 288433 | 21330 | 8599 | 10638 | 57686700 | 0.08 | 0.06 | 0.06 | 21.004 |
| P21 | M | PL | 308316 | 24059 | 11403 | 9950 | 61663300 | 0.08 | 0.06 | 0.06 | 28.425 |
| P21 | F | PL | 345480 | 20360 | 10225 | 9694 | 69095900 | 0.08 | 0.06 | 0.06 | 25.668 |
| P21 | F | PL | 332853 | 21284 | 10112 | 8311 | 66570700 | 0.07 | 0.06 | 0.06 | 23 |
| P21 | F | PL | 290021 | 22434 | 12493 | 9411 | 58004100 | 0.08 | 0.06 | 0.07 | 30.701 |
| P21 | M | PL | 331717 | 20207 | 9143 | 8161 | 66343400 | 0.08 | 0.06 | 0.07 | 24.881 |
| P28 | F | PL | 334221 | 21149 | 9340 | 3382 | 66844100 | 0.08 | 0.06 | 0.07 | 24.111 |
| P28 | M | PL | 349135 | 24051 | 10088 | 3064 | 69827000 | 0.07 | 0.06 | 0.07 | 24.151 |
| P28 | F | PL | 315814 | 20152 | 9466 | 3894 | 63162800 | 0.08 | 0.06 | 0.07 | 24.348 |
| P28 | M | PL | 327118 | 22676 | 10594 | 2998 | 65423600 | 0.07 | 0.06 | 0.07 | 23.836 |
| P28 | F | PL | 281363 | 20116 | 8572 | 3229 | 56272600 | 0.08 | 0.06 | 0.08 | 26.878 |
| P35 | F | PL | 310024 | 21903 | 11966 | 3315 | 62004700 | 0.08 | 0.06 | 0.08 | 29.504 |
| P35 | M | PL | 316646 | 24474 | 11496 | 3703 | 63329200 | 0.08 | 0.06 | 0.08 | 31.391 |
| P35 | F | PL | 348477 | 20384 | 11580 | 3961 | 69695400 | 0.08 | 0.06 | 0.07 | 25.467 |
| P35 | M | PL | 292859 | 23607 | 10447 | 3371 | 58571800 | 0.08 | 0.06 | 0.08 | 30.442 |
| P35 | F | PL | 269859 | 18259 | 8197 | 3126 | 53971800 | 0.08 | 0.06 | 0.08 | 23.996 |
| P60 | M | PL | 241252 | 19163 | 9924 | 1754 | 48250300 | 0.09 | 0.06 | 0.09 | 28.704 |
| P60 | F | PL | 350853 | 20230 | 9237 | 1670 | 70170600 | 0.08 | 0.06 | 0.09 | 24.546 |
| P60 | F | PL | 260818 | 20098 | 10681 | 2663 | 52163600 | 0.09 | 0.07 | 0.08 | 35.123 |
| P60 | M | PL | 285108 | 19769 | 10217 | 2218 | 57021500 | 0.08 | 0.06 | 0.08 | 28.221 |
| P60 | F | PL | 286159 | 20608 | 9178 | 1948 | 57231900 | 0.08 | 0.06 | 0.08 | 24.722 |
| P7 | F | vEN | 1629750 | 42625 | 19685 | 2912 | 325951000 | 0.07 | 0.05 | 0.06 | 18.961 |
| P7 | M | vEN | 1468690 | 46019 | 15240 | 2025 | 293739000 | 0.06 | 0.06 | 0.07 | 18.215 |
| P7 | M | vEN | 1558320 | 46754 | 17412 | 2265 | 311663000 | 0.06 | 0.05 | 0.07 | 18.951 |
| P7 | M | vEN | 1304060 | 47149 | 17602 | 2346 | 260811000 | 0.06 | 0.05 | 0.07 | 18.975 |
| P7 | F | vEN | 1698750 | 46307 | 18003 | 3154 | 339751000 | 0.06 | 0.05 | 0.06 | 18.217 |
| P14 | M | vEN | 1471930 | 43938 | 17491 | 1315 | 294386000 | 0.07 | 0.08 | 0.09 | 18.533 |
| P14 | F | vEN | 1748400 | 40864 | 17522 | 1503 | 349680000 | 0.07 | 0.07 | 0.09 | 18.986 |
| P14 | M | vEN | 1696020 | 48014 | 21096 | 1556 | 339203000 | 0.07 | 0.07 | 0.09 | 21.402 |
| P14 | F | vEN | 1561710 | 47578 | 18963 | 1750 | 312341000 | 0.07 | 0.08 | 0.09 | 21.002 |
| P14 | F | vEN | 1456410 | 46760 | 16702 | 1641 | 291283000 | 0.07 | 0.08 | 0.09 | 21.044 |
| P21 | M | vEN | 1639490 | 60265 | 21702 | 7097 | 327898000 | 0.07 | 0.05 | 0.08 | 29.189 |
| P21 | F | vEN | 1860210 | 52548 | 16403 | 5588 | 372043000 | 0.07 | 0.06 | 0.09 | 25.801 |
| P21 | F | vEN | 1959740 | 59396 | 20856 | 5247 | 391947000 | 0.06 | 0.05 | 0.08 | 22.891 |
| P21 | F | vEN | 1387660 | 47494 | 17090 | 6416 | 277532000 | 0.07 | 0.05 | 0.08 | 27.328 |
| P21 | M | vEN | 1779930 | 56518 | 18697 | 7278 | 355986000 | 0.06 | 0.05 | 0.07 | 25.102 |
| P28 | F | vEN | 1784300 | 50095 | 26338 | 3232 | 356860000 | 0.06 | 0.06 | 0.08 | 24.366 |
| P28 | M | vEN | 1852680 | 56981 | 24582 | 2883 | 370536000 | 0.06 | 0.07 | 0.09 | 24.08 |
| P28 | F | vEN | 1786260 | 56952 | 24338 | 5544 | 357252000 | 0.06 | 0.07 | 0.06 | 24.777 |
| P28 | M | vEN | 2050660 | 58691 | 26887 | 3588 | 410133000 | 0.06 | 0.07 | 0.08 | 24.655 |
| P28 | F | vEN | 1834560 | 55155 | 22372 | 4278 | 366912000 | 0.07 | 0.07 | 0.08 | 27.564 |
| P35 | F | vEN | 1693650 | 52208 | 25760 | 4441 | 338730000 | 0.07 | 0.06 | 0.08 | 28.801 |
| P35 | M | vEN | 2008970 | 58939 | 23041 | 3582 | 401794000 | 0.06 | 0.06 | 0.08 | 25.818 |
| P35 | F | vEN | 1758030 | 48912 | 21599 | 4160 | 351605000 | 0.07 | 0.06 | 0.07 | 24.992 |
| P35 | M | vEN | 1820010 | 59706 | 26167 | 3772 | 364002000 | 0.07 | 0.06 | 0.08 | 27.482 |
| P35 | F | vEN | 1907040 | 58966 | 20316 | 3170 | 381408000 | 0.06 | 0.06 | 0.08 | 23.836 |
| P60 | M | vEN | 1663080 | 55057 | 25654 | 2405 | 332616000 | 0.08 | 0.07 | 0.08 | 28.374 |
| P60 | F | vEN | 2012800 | 53278 | 21525 | 2745 | 402560000 | 0.07 | 0.06 | 0.07 | 25.424 |
| P60 | F | vEN | 1949860 | 61929 | 26338 | 2034 | 389971000 | 0.08 | 0.07 | 0.09 | 26.791 |
| P60 | M | vEN | 1958370 | 62065 | 27606 | 2371 | 391675000 | 0.07 | 0.06 | 0.08 | 24.762 |
| P60 | F | vEN | 1882470 | 54329 | 24810 | 2600 | 376495000 | 0.08 | 0.07 | 0.07 | 24.882 |
