## Supplementary Table 3 for "Mouse paralaminar amygdala excitatory neurons migrate and mature during adolescence"

**Supplementary Table 3: Human Case Table**

| Case no. | Age | Gender | PMI | Experimental Use | Neuropathology diagnosis | Clinical history |
|---|---|---|---|---|---|---|
| 1 | 18 GW | M | 24h | IHC | control | prematurity |
| 2 | 22 GW | M | 24h | IHC | control | urethral stenosis |
| 3 | 22 GW | M | 48h | IHC | control | spontaneous abortion |
| 4 | 27 GW | M | 12h | IHC | control | intrauterine growth retardation |
| 5 | 28 GW | M | 24h | IHC | control | prematurity |
| 6 | 29 GW | F | 24h | IHC | control | Necrotizing enterocolitis |
| 7 | 37 GW | M | 48h | IHC | control | VATER malformation |
| 8 | 38 GW | M | 50h | IHC | control | bronchopulmonary dysplasia |
| 9 | term | M | 36h | IHC | control | Intra-partem death, presumed nuchal cord |
| 10 | term | M | 14h | IHC | control | chondrodysplasia |
| 11 | term | M | 11h | IHC | control | cardiac dysfunction |
| 12 | term | M | 24h | IHC | control | hydronephrosis |
| 13 | 1 day | F | 25h | IHC | control | renal hypoplasia; hypoplastic lung |
| 14 | 2 day | M | 36h | IHC | control | hypoplastic L heart |
| 15 | 3 months | M | 20h | IHC | control | diaphragmatic hernia |
| 16 | 7 months | M | 14h | IHC | control | Tetralogy of Fallot |
| 17 | 11 months | M | 21h | IHC | control | bacterial pneumonia |
| 18 | 22 months | M | 38h | IHC | control | cardiovascular failure |
| 19 | 2 years | M | 19h | IHC | control | leukemia |
| 20 | 3 years | M | 25h | IHC | control | esophageal atresia, tracheoesophageal fistula |
| 21 | 13 years | M | 12h | IHC | control | focal segmental glomerulosclerosis |
