## Supplementary Table 4 for "Mouse paralaminar amygdala excitatory neurons migrate and mature during adolescence"

| Supplementary Table 4: Antibodies | | | | |
|---|---|---|---|---|
| **Antigen** | **Species** | **Dilution** | **Manufacturer** | **Cat. No.** |
| APC | Mouse | 1:1000 | Millipore-Sigma | OP80 |
| BrdU | Rat | 1:200 | Abcam | ab6326 |
| Cleaved Caspase-3 | Rabbit | 1:500 | Cell Signal | 9664S |
| CoupTFII | Rabbit | 1:500 | Abcam | AB211777 |
| CoupTFII | Mouse | 1:250 IHC 1: 100 EM | R&D Systems | PP-H7147-00 |
| Ctip2 | Rat | 1:200 | Abcam | ab18465 |
| Dcx | Rabbit | 1:500 IHC 1: 100 EM | Cell Signal | 4604 |
| Dcx | Guinea pig | 1:500 | EMD Millipore | AB2253 |
| FoxP2 | Rabbit | 1:500 | Atlas Antibodies | hpa000382 |
| FoxP2 | Goat | 1:500 | Santa Cruz Biotechnology | 517261 |
| GFAP | Rabbit | 1:500 | EMD Millipore | Z0334 |
| Mcherry (tdTomato) | Rat | 1:200 | Invitrogen | M11217 |
| NeuN | Chicken | 1:200 | EMD Millipore | ABn91 |
| Olig2 | Rabbit | 1:500 | EMD Millipore | AB9610 |
| Pax6 | Rabbit | 1:500 | Bio-Legend | 901301 |
| Psa-Ncam | Mouse | 1:1000 | EMD Millipore | MAB5324 |
| SatB2 | Rabbit | 1:500 | Abcam | AB34735 |
| Sp8 | Goat | 1:500 | Santa Cruz Biotechnology | SC104664 |
| Tbr1 | Chicken | 1:500 IHC 1: 100 EM | EMD Millipore | AB2261 |
| Tbr1 | Rabbit | 1:500 | EMD Millipore | AB10554 |
| Alexa Fluor® 488 Donkey anti-Rabbit IgG (H+L) | Donkey | 1:2000 | Invitrogen | A32790 |
| Alexa Fluor® 555 Donkey anti-Rabbit IgG (H+L) | Donkey | 1:2000 | Invitrogen | A32794 |
| Donkey anti-Rabbit IgG (H+L) Highly Cross-Adsorbed Secondary Antibody, Alexa Fluor™ Plus 647 | Donkey | 1:2000 | Invitrogen | a32795 |
| Goat anti-Rabbit IgG (H+L) Cross-Adsorbed Secondary Antibody, Alexa Fluor™ 647 | Goat | 1:2000 | Invitrogen | A21244 |
| Goat anti-Chicken IgY (H+L) Cross-Adsorbed Secondary Antibody, Alexa Fluor™ Plus 488 | Goat | 1:2000 | Invitrogen | A32931 |
| Alexa Fluor® 488 AffiniPure Donkey Anti-Chicken IgY (IgG) (H+L) | Donkey | 1:2000 | Jackson ImmunoResearch | 703-545-155 |
| Alexa Fluor® Cy3 AffiniPure Donkey Anti-Chicken IgY (IgG) (H+L) | Donkey | 1:2000 | Jackson ImmunoResearch | 703-165-155 |
| Alexa Fluor® 647 AffiniPure Donkey Anti-Chicken IgY (IgG) (H+L) | Donkey | 1:2000 | Jackson ImmunoResearch | 703-605-155 |
| Alexa Fluor® 488 AffiniPure Goat Anti-Rat IgG (H+L) | Goat | 1:2000 | Jackson ImmunoResearch | 112-545-167 |
| Cy™3 AffiniPure Donkey Anti-Rat IgG (H+L) | Donkey | 1:2000 | Jackson ImmunoResearch | 712-165-153 |
| Alexa Fluor® 647 AffiniPure Donkey Anti-Rat IgG (H+L) | Donkey | 1:2000 | Jackson ImmunoResearch | 712-605-153 |
| Alexa Fluor® 555 Goat anti-Guinea Pig IgG (H+L) | Goat | 1:2000 | Invitrogen | A21435 |
| Alexa Fluor® 647 AffiniPure Donkey Anti-Guinea Pig IgG (H+L) | Donkey | 1:2000 | Jackson ImmunoResearch | 706-605-148 |
| Goat anti-Guinea Pig IgG (H+L) Highly Cross-Adsorbed Secondary Antibody, Alexa Fluor™ 647 | Goat | 1:2000 | Invitrogen | A21450 |
| Alexa Fluor® 488 AffiniPure Donkey Anti-Goat IgG (H+L) | Donkey | 1:2000 | Jackson ImmunoResearch | 705-545-003 |
| Alexa Fluor® 488 AffiniPure Donkey Anti-Mouse IgG (H+L) | Donkey | 1:2000 | Invitrogen | A32766 |
| Alexa Fluor® 488 AffiniPure Goat Anti-Mouse IgM (H+L) | Goat | 1:2000 | Invitrogen | A21042 |
| Alexa Fluor® 555 AffiniPure Donkey Anti-Mouse IgG (H+L) | Donkey | 1:2000 | Invitrogen | A32773 |
| Alexa Fluor® 555 AffiniPure Goat Anti-Mouse IgM (H+L) | Goat | 1:2000 | Invitrogen | A21426 |
| Donkey anti-Mouse IgG (H+L) Highly Cross-Adsorbed Secondary Antibody, Alexa Fluor™ Plus 647 | Donkey | 1:2000 | Invitrogen | A32787 |
| Alexa Fluor® 647 AffiniPure Goat Anti-Mouse IgM (H+L) | Goat | 1:2000 | Invitrogen | A21238 |

| |
|---|
| **Lot No(s).** |
| 3307277 |
| GR3365969-10 |
| 22 |
| GR3396420-9 |
| A-2 |
| gr243609-4 |
| 6, 7 |
| 3847922<br>3777998<br>3601335 |
| B115858 |
| G2216 |
| 3194598 |
| WI334529<br>XC345714 |
| 3788621<br>3788621 |
| 3755328 |
| B201255 |
| 3607061 |
| GR3354380-1 |
| G0516 |
| 3598056<br>3105253<br>3811721 |
| 3598056<br>3105253<br>3811721 |
| TI271741 |
| WG322207 |
| UG289708 |
| 1463166 |
| TI271747 |
| 146581 |
| 144929 |
| 153969 |
| 148029 |
| 156617 |
| 154486 |
| 2076358 |
| 149191 |
| 2110845 |
| 148783 |
| TI271738 |
| 2079371 |
| TI271028 |
| 2001027 |
| TJ271040 |
| 2112240 |
